# RCGAToolbox: A real-coded genetic algorithm software for parameter estimation of kinetic models

**DOI:** 10.1101/2021.02.15.431062

**Authors:** Kazuhiro Maeda, Fred C. Boogerd, Hiroyuki Kurata

## Abstract

**Summary:** Kinetic modeling is essential in understanding the dynamic behavior of biochemical networks, such as metabolic and signal transduction pathways. However, parameter estimation remains a major bottleneck in the development of kinetic models. We present RCGAToolbox, software for real-coded genetic algorithms (RCGAs), which accelerates the parameter estimation of kinetic models. RCGAToolbox provides two RCGAs: the unimodal normal distribution crossover with minimal generation gap (UNDX/MGG) and real-coded ensemble crossover star with just generation gap (REX^star^/JGG), using the stochastic ranking method. The RCGAToolbox also provides user-friendly graphical user interfaces.

**Availability and implementation:** RCGAToolbox is available from https://github.com/kmaeda16/RCGAToolbox under GNU GPLv3, with application examples. The user guide is provided in the Supplementary Material. RCGAToolbox runs on MATLAB in Windows, Linux, and macOS.

**Supplementary information:** Supplementary Material is available at *Bioinformatics* online.

## 1. Introduction

Kinetic modeling and simulations are essential for understanding complex biological systems (Kitano, 2002). Kinetic models consist of sets of differential equations with various model parameters. As most model parameters have not been measured experimentally, they need to be estimated to best fit experimental training data. Parameter estimation remains a major bottleneck in the modeling process owing to the large number of model parameters, nonlinear dynamics, and local optima (Banga, 2008). Several related software tools have been developed, for example, AMIGO2 (Balsa-Canto, et al., 2016), MEIGO (Egea, et al., 2014), CADLIVE Toolbox (Inoue, et al., 2014), and libRGCA (Maeda, et al., 2018). Such tools are meant to be used by experts, as the lack of user-friendliness hampers general usage by capable yet inexperienced scientists. Therefore, we introduced RCGAToolbox. RCGAToolbox is a parameter estimation software that provides real-coded genetic algorithms (RCGAs). It implements two proven RCGAs: the unimodal normal distribution crossover with minimal generation gap (UNDX/MGG) (Ono and Kobayashi, 1997; Satoh, et al., 1997) and the real-coded ensemble crossover star with just generation gap (REX^star^/JGG) (Kobayashi, 2009). The toolbox provides easy-to-use graphical user interfaces (GUIs) to perform these RCGAs.

## 2. Features

RCGAToolbox is a MATLAB-based multiplatform application for parameter estimation of kinetic models. For use of RCGAs, installation of any other toolbox is not required. However, to perform import/export of SBML models, fast simulation, local search methods, and parallelization, the corresponding third-party toolboxes need to be installed additionally. The user guide is provided as Supplementary Material with step-by-step examples. The GUIs allow users to perform straightforward parameter estimation and generate MATLAB scripts for advanced use of RCGAToolbox.

### 2.1. Problem definition

Kinetic models include various parameters, for instance maximum velocity (V_max_) and Michaelis constant (K_m_). Parameter estimation minimizes the objective function for a specific model. The objective function is the error between the model prediction and experimental training data. Parameter estimation often involves algebraic or dynamic constraints based on prior knowledge or an educated guess, leading to a constrained parameter estimation problem.

### 2.2. Numerical methods

RCGAs are population-based stochastic optimization algorithms. Initial populations are randomly generated in which each individual is characterized by a set of different values for model parameters. By repeating selection and reproduction operations, RCGAs evolve the population and eventually obtain the individuals (the sets of model parameters) providing an optimal fit for experimental data. The two RCGAs, UNDX/MGG and REX^star^/JGG, have been employed previously for parameter estimation (Jahan, et al., 2016; Maeda, et al., 2019). UNDX/MGG prevents the population from being trapped in local optima by allowing only small changes per generation, whereas REX^star^/JGG evaluates the search space and moves the population in a favorable direction.

RCGAToolbox provides the stochastic ranking method (Runarsson and Yao, 2000) for efficient parameter estimation with constraints. Local optimizers can be combined with RCGAs to rapidly refine solutions. Parallel computation is supported, which greatly accelerates parameter estimation (see **Section 7.5** of Supplementary Material). For further acceleration, RCGAToolbox provides convenient access to C-based fast ordinary differential equation solvers provided by third-parties (see **Section 7.1** of Supplementary Material). RCGAToolbox can import Systems Biology Markup Language (SBML) or work with arbitrary user-defined models (see **Section 5.1** of Supplementary Material). The import and export of SBML models are realized by IQM Tools (IntiQuan), enabling conversion between SBML models and MATLAB structures. RCGAToolbox is flexible and extensible and can handle arbitrary objective functions. Users can also quickly implement their evolutionary algorithms by modifying template codes (see **Section 5.3** of Supplementary Material).

## 3. Performance

We performed benchmark experiments for the RCGAs provided in RCGAToolbox. Altogether, we tested 33 problems: 27 mathematical benchmarks and 6 parameter estimations. The performance of REX^star^/JGG from RCGAToolbox was comparable to that of eSS [R2019 version provided by MEIGO (Egea, et al., 2014)] (see **Section 7.2-7.4** of Supplementary Material). We also found that REX^star^/JGG outperformed eSS in one of the parameter estimation problems. The source codes for the computational experiments are included in the distribution of RCGAToolbox.

## Availability and requirements

Project name: RCGAToolbox

Project home page: https://github.com/kmaeda16/RCGAToolbox

Operating system(s): Platform independent

Programming language: MATLAB

Other requirements: Windows, Linux, macOS with MATLAB R2016a or later

License: GNU GPL v3

Any restrictions to use by non-academics: License needed

## Declarations

### Availability of data and materials

The data underlying this article will be shared on reasonable request to the corresponding author.

## Acknowledgements

The authors thank Editage for reviewing and editing the manuscript.

## Authors’ contributions

With support and guidance from FCB and HK, KM developed the software and performed the computational experiments. KM, FCB, and HK analyzed the data and wrote the manuscript.

