## Supplementary Material for "RCGAToolbox: A real-coded genetic algorithm software for parameter estimation of kinetic models"

### RCGAToolbox User Guide

### 1. Introduction

#### 1.1. What is RCGAToolbox?

RCGAToolbox is a MATLAB toolbox that contains two real-coded genetic algorithms (RCGAs): the unimodal normal distribution crossover with minimal generation gap (UNDX/MGG) and the real-coded ensemble crossover star with just generation gap ($\text{REX}^{\text{star}}$/JGG). The stochastic ranking method is implemented to efficiently handle constrained optimization problems. RCGAToolbox not only provides access to RCGAs but also several useful features for parameter estimation in systems biology. RCGAToolbox is licensed under [the GNU General Public License v3.0].

This document includes contents of our previous paper (Maeda, et al., 2018) with permission from the publisher (Information Processing Society of Japan), including descriptions of RCGAs (**Section 6** and **Figure 5**) and test problems (**Tables 5–7**, **Figures 9–10**, and **Appendices C–E**).

#### 1.2. Problem formulation

RCGAs in RCGAToolbox handle the following constrained optimization problem.

$$\text{Minimize} \quad f(\mathbf{x}), \tag{1a}$$

$$\text{Subject to} \quad \mathbf{g}(\mathbf{x}) \le \mathbf{0}, \tag{1b}$$

$$\mathbf{x}^{lb} \le \mathbf{x} \le \mathbf{x}^{ub}, \tag{1c}$$

where $\mathbf{x} = (x_1, x_2, \ldots)$ is the vector with decision variables, $f$ is the objective function, and $\mathbf{g} = (g_1, g_2, \ldots)$ is the vector with constraint functions. A positive value of $g_i$ indicates that the $i^{\text{th}}$ constraint is violated. $\mathbf{x}^{lb}$ and $\mathbf{x}^{ub}$ are the lower and upper bounds, respectively. Eq. (1c) defines the search space. Eqs. (1a-c) together represent a constrained optimization problem. In an *unconstrained* optimization problem, Eq. (1b) is omitted.

In the context of parameter estimation, the decision variables ($\mathbf{x}$) indicate kinetic parameters to be estimated, and $f$ evaluates the model's fit to experimental data, whereas $g_i$'s are constraint functions that the kinetic model must satisfy.

### 2. Installation

#### 2.1. Requirements

- MATLAB R2016a or later. We confirmed that RCGAToolbox runs on (i) Windows 10 (2004) with MATLAB R2016a, (ii) SUSE Linux Enterprise Server 11 (x86_64) with MATLAB R2016a, and (iii) macOS Big Sur (11.1, Intel CPU) with MATLAB R2020b.
- Optional requirements
  - ✧ Parallel Computing Toolbox is required for parallel computation (opts.n_par > 1, see below). It is not required for sequential computation.
  - ✧ Optimization Toolbox is required for local optimization using fmincon (opts.local = 1, see below). It is not required if the local optimization function is not used.
  - ✧ IQM Tools [formerly known as SBToolbox2/SBPD (Schmidt, 2007; Schmidt and Jirstrand, 2006)] are required for handling Systems Biology Markup Language (SBML) and a fast simulation (fast_flag = 2).
  - ✧ SundialsTB is required for a fast simulation with CVODE (fast_flag = 1).

For installation of IQM Tools and SundialsTB, see **Appendix A**. For the list of functions provided by RCGAToolbox, see **Appendix B**.

#### 2.2. Installation

1. Download RCGAToolbox from https://github.com/kmaeda16/RCGAToolbox.
2. Place the directory RCGAToolbox somewhere favorable (e.g. Documents/MATLAB/).
3. Run the installation script **RCGAToolbox_Install.m** under the directory RCGAToolbox/install/.

#### 2.3. Uninstallation

1. Run the uninstallation script **RCGAToolbox_Uninstall.m** under the directory RCGAToolbox/install/.
2. Delete the directory RCGAToolbox.

#### 2.4. Diagnosis

RCGAToolbox/install/**RCGAToolbox_Diagnosis.m** is the self-diagnosis script that checks whether the RCGAToolbox is properly installed. It also tests the RCGAToolbox functions that depend on optional toolboxes. For the diagnosis, run **RCGAToolbox_Diagnosis.m** under the directory RCGAToolbox/install/.

### 3. Quick start

#### 3.1. General optimization problem

RCGAToolbox/doc/demo/general/**run_QuickStart.m** demonstrates how to run RCGAs using RCGAToolbox. Simply define your problem and assign it to either function **RCGA_UNDXMGG** or **RCGA_REXstarJGG** to solve.

RCGAToolbox/doc/demo/general/run_QuickStart.m

```
% This script demonstrates how to run a real-coded genetic algorithm to
% solve an example constrained optimization problem.
%
% --------------------- Example Problem ---------------------
% Minimize:
%   f = x(1)^2 + x(2)^2 + ... + x(10)^2
%
% Subject to:
%   g(1) = x(1) * x(2) + 1 <= 0
%   g(2) = x(1) + x(2) + 1 <= 0
%   -5.12 <= x(i) <= 5.12 for all i
%
%
% Global minimum is f = 3, g(1) = 0, g(2) = 0 at x = (-1.618, 0.6180, 0, 0,
% 0, 0, 0, 0, 0, 0) or at x = (0.6180, -1.618, 0, 0, 0, 0, 0, 0, 0, 0)
% -----------------------------------------------------------

clearvars;

% ========= Problem Settings ========= %
problem.n_gene = 10; % Number of Decision Variables
problem.n_constraint = 2; % Number of Constraints
problem.fitnessfun = @Fitness_Example; % Fitness Function
problem.decodingfun = @Decoding_Example; % Decoding Function

% ========== Executing RCGA ========== %
% Results = RCGA_UNDXMGG(problem); % UNDX/MGG
Results = RCGA_REXstarJGG(problem); % REXstar/JGG
```

Executing **run_QuickStart.m** gives you the following messages in the MATLAB Command Window.

```
>> run_QuickStart

================================================

          RCGA_REXstarJGG by RCGAToolbox

================================================

------------------- Problem --------------------
         n_gene :  10
    n_constraint :  2
      fitnessfun :  Fitness_Example
     decodingfun :  Decoding_Example
------------------- Options --------------------
    n_population :  300
      n_children :  300
        n_parent :  11
       t_rexstar :  6
  selection_type :  0
              Pf :  0.45
           local :  0
          maxgen :  1000
         maxtime :  600
         maxeval :  Inf
             vtr :  -Inf
           n_par :  1
    output_intvl :  1
  out_transition :  None
        out_best :  None
  out_population :  None
      out_report :  None
interimreportfun :  RCGAdefaultinterimreportfun
  finalreportfun :  RCGAdefaultfinalreportfun
------------------------------------------------

```

```
RCGA_REXstarJGG started.
Elapsed Time = 5.748680e-02, Generation = 1, f = 4.451477e+01, phi = 0.000000e+00
Elapsed Time = 1.497601e-01, Generation = 2, f = 2.558670e+01, phi = 0.000000e+00
Elapsed Time = 2.270435e-01, Generation = 3, f = 1.315321e+01, phi = 0.000000e+00
...
Elapsed Time = 3.363103e+01, Generation = 998, f = 3.142870e+00, phi = 0.000000e+00
Elapsed Time = 3.366195e+01, Generation = 999, f = 3.142870e+00, phi = 0.000000e+00
Elapsed Time = 3.369350e+01, Generation = 1000, f = 3.142870e+00, phi = 0.000000e+00
Maximum number of generations (maxgen) reached.
```

In the messages, $f$ indicates the objective function value, and *phi* indicates the penalty function value given as $\phi(\mathbf{x}) = \sum_{i=1}^{m} \left[ \max\left\{0, g_i(\mathbf{x})\right\} \right]^2$ (see **Section 1.2** for $g_i$ and **x**). $m$ is the number of constraint functions.

As shown in **run_QuickStart.m**, the following four fields of the structure **problem** need to be set before it can be passed to **RCGA_UNDXMGG** or **RCGA_REXstarJGG**:

- **problem.n_gene**: The number of decision variables.
- **problem.n_constraint**: The number of constraint functions. For unconstrained problems, it is zero.
- **problem.fitnessfun**: The function handle for a fitness function that takes **x** as an input and returns $f$ and **g**. For unconstrained problems, it returns only $f$.
- **problem.decodingfun**: The function handle for a decoding function that defines the search space and how to decode genes to **x**.

In **run_QuickStart.m**, an RCGA solves the following constrained optimization problem:

$$\text{Minimize} \quad f(\mathbf{x}) = \sum_{i=1}^{10} x_i^2 , \tag{2a}$$

$$\text{Subject to} \quad g_1(\mathbf{x}) = x_1 x_2 + 1 \le 0 , \tag{2b}$$

$$g_2(\mathbf{x}) = x_1 + x_2 + 1 \le 0 , \tag{2c}$$

$$-5.12 \le x_i \le 5.12 , \tag{2d}$$

where $f$ is the objective function and $g_i$ is the $i^{\text{th}}$ constraint function. **x** is a decision variable vector: **x** = $(x_1, x_2, ..., x_{10})$. The fitness function and the decoding function for this constrained optimization problem are given as **Fitness_Example.m** and **Decoding_Example.m**, respectively:

RCGAToolbox/doc/demo/general/Fitness_Example.m

```
function [f, g] = Fitness_Example(x)
% Fitness_Example is an example of fitness function.
%
% [SYNTAX]
% [f, g] = Fitness_Example(x)
%
% [INPUT]
% x : Decision variables.
%
% [OUTPUT]
% f : Objective function to be minimized.
% g : Constraint functions to be less than zero.
%
% --------------------- Example Problem ---------------------
% Minimize:
%   f = x(1)^2 + x(2)^2 + ... + x(10)^2
%
% Subject to:
%   g(1) = x(1) * x(2) + 1 <= 0
%   g(2) = x(1) + x(2) + 1 <= 0
%   -5.12 <= x(i) <= 5.12 for all i
%
%
% Global minimum is f = 3, g(1) = 0, g(2) = 0 at x = (-1.618, 0.6180, 0, 0,
% 0, 0, 0, 0, 0, 0) or at x = (0.6180, -1.618, 0, 0, 0, 0, 0, 0, 0, 0)
% ----------------------------------------------------------

f    = sum( x .^ 2 );

g(1) = x(1) * x(2) + 1;
g(2) = x(1) + x(2) + 1;
```

RCGAToolbox/doc/demo/general/Decoding_Example.m

```
function x = Decoding_Example(gene)
% Decoding_Example is an example of decoding function for
% "Fitness_Example.m". "gene" takes values from 0 to 1. The purpose of
% decoding functions is to change the value range, i.e. to decode "gene"
% and return it as x.
%
% [SYNTAX]
% x = Decoding_Example(gene)
%
% [INPUT]
% gene : Encoded decision variables.
%
% [OUTPUT]
% x    : Decoded decision variables.
%
% --------------------- Example Problem ---------------------
% Minimize:
%   f = x(1)^2 + x(2)^2 + ... + x(10)^2
%
% Subject to:
%   g(1) = x(1) * x(2) + 1 <= 0
%   g(2) = x(1) + x(2) + 1 <= 0
%   -5.12 <= x(i) <= 5.12 for all i
%
%
% Global minimum is f = 3, g(1) = 0, g(2) = 0 at x = (-1.618, 0.6180, 0, 0,
% 0, 0, 0, 0, 0, 0) or at x = (0.6180, -1.618, 0, 0, 0, 0, 0, 0, 0, 0)
% -----------------------------------------------------------

lb = -5.12;
ub =  5.12;

x = gene * ( ub - lb ) + lb;
```

Note that for *unconstrained* optimization problems, fitness functions should return only *f*. In RCGAToolbox, decision variables are internally expressed as a vector with a range of 0–1. The internal decision variable is called a **gene**. Decoding functions take genes as input, decode these to decision variables **x**, and return these. Decoding functions provide flexibility in defining search space: either linear, logarithmic, continuous, or discontinuous.

The functions **RCGA_UNDXMGG** and **RCGA_REXstarJGG** return the structure **Results**, which has four fields: **Transition**, **Best**, **FinalPopulation**, and **end_crit**. The former three has the same information as the transition file, the best individual file, and the final population file, respectively (see **Section 4.1.1** for these files). **end_crit** is the exit flag, whereas **end_crit** = 0 indicates that the RCGAs successfully found a solution. A solution is the decision variable vector that provides an objective function value (*f*) less than a user-defined value (called the value to be reached or **vtr**) with all the constraints satisfied ($\mathbf{g}_i \leq 0$). Non-zero **end_crit** indicates the RCGAs finished without finding a solution: **end_crit** = 1, 2, or 3 indicates "the maximal number of generations (maxgen) was reached," "the maximum CPU time (maxtime) was reached," or "the maximum number of fitnessfun evaluations (maxeval) was reached," respectively.

##### 3.2. Parameter estimation in systems biology

An example in this section requires IQM Tools. If you have not installed IQM Tools, please use sample scripts in RCGAToolbox/doc/demo/*PE_woIQM*/ instead of RCGAToolbox/doc/demo/*PE*/.

RCGAToolbox/doc/demo/PE/**run_QuickStart.m** demonstrates how to use RCGAs for a parameter estimation problem in systems biology. Simply call **RCGA_UNDXMGG_PE** or **RCGA_REXstarJGG_PE** with a kinetic model (**modelfile**), target experimental data (**measurement**), and a decoding function (**decodingfun**).

**Model_Example_odefun.m** is an ODE file (IQM Tools format) representing a simple kinetic model with two state variables and seven model parameters. **Measurement_Example.csv** is a CSV file that represents the target experimental data. **Decoding_Example.m** is the decoding function.

RCGAToolbox/doc/demo/PE/run_QuickStart.m

```
% This script demonstrates how to run a real-coded genetic algorithm to
% estimate model parameters in an example kinetic model.
%
% ------------------------ Example Kinetic Model ------------------------
% - INITIAL CONDITION
% X1 = 0
% X2 = 0
%
% - PARAMETERS
% X0 = 0.1
% k1 = 1
% k2 = 1
% k3 = 1
% K2 = 1
% K3 = 1
% rootCompartment = 1
%
% - VARIABLES
% X12 = X1 + X2
%
% - REACTIONS
% v1 = k1 * X0
% v2 = k2 * (X1/rootCompartment) / (K2 + (X1/rootCompartment))
% v3 = k3 * (X2/rootCompartment) / (K3 + (X2/rootCompartment))
%
% - BALANCE
% X1_dot = v1 - v2;
% X2_dot = v2 - v3;
% -----------------------------------------------------------------------

clear mex;
clear all;
close all;

% ========= Problem Settings ========= %
```

```
modelfile = @Model_Example_odefun; % ODE file (IQM Tools format)
decodingfun = @Decoding_Example; % Decoding Function
% measurement = 'Measurement_Example.xls'; % Measurement File (EXCEL format)
measurement = 'Measurement_Example.csv'; % Measurement File (CSV)

% ========== Executing RCGA ========== %
% Results = RCGA_UNDXMGG_PE(modelfile,decodingfun,measurement); % UNDX/MGG
Results = RCGA_REXstarJGG_PE(modelfile,decodingfun,measurement); % REXstar/JGG
```

Instead of **RCGA_REXstarJGG** (which is used in **Section 3.1**), **RCGA_REXstarJGG_PE** is used here, where "PE" stands for parameter estimation. Executing **run_QuickStart.m** provides the following messages in the MATLAB Command Window. This calculation takes 10 min to complete. To abort the running script, press 'Control + C' in the MATLAB Command Window.

```
>> run_QuickStart
Reading Measurement_Example.csv ... Finished.

================================================

          RCGA_REXstarJGG by RCGAToolbox

================================================

------------------- Problem --------------------
         n_gene :  7
   n_constraint :  0
     fitnessfun :  @(x)fitnessfun(x,Simulation,model,mst,simopts)
    decodingfun :  Decoding_Example
------------------- Options --------------------
   n_population :  300
     n_children :  300
       n_parent :  8
      t_rexstar :  6
 selection_type :  0
             Pf :  0.45
          local :  0
```

```
          maxgen :  1000
         maxtime :  600
         maxeval :  Inf
             vtr :  -Inf
           n_par :  1
    output_intvl :  1
  out_transition :  None
        out_best :  None
  out_population :  None
      out_report :  None
interimreportfun                                                   :
@(elapsedTime,generation,problem,opts,Population,best)interimreportfun(elapsedTi
me,generation,problem,opts,Population,best,Simulation,model,mst,simopts)
  finalreportfun :  RCGAdefaultfinalreportfun
------------------------------------------------

RCGA_REXstarJGG started.
Elapsed Time = 2.339310e+00, Generation = 1, f = 6.562991e-03
Elapsed Time = 5.673242e+00, Generation = 2, f = 3.941497e-03
Elapsed Time = 7.961011e+00, Generation = 3, f = 2.229762e-03
...
Elapsed Time = 5.967471e+02, Generation = 290, f = 1.746201e-06
Elapsed Time = 5.987983e+02, Generation = 291, f = 1.746201e-06
Elapsed Time = 6.008096e+02, Generation = 292, f = 1.746201e-06
Maximum CPU time (maxtime) reached.

--- Best parameter set (f = 1.746201e-06) ---
X0 = 2.358726e-01
k1 = 4.199921e-01
k2 = 1.329753e+00
k3 = 1.318928e+00
K2 = 1.168590e+00
K3 = 1.153963e+00
rootCompartment = 1.179145e+00
```

In the command window, *f* is the objective function value, which is defined as the squared difference between model predictions and experimental data: $f(\mathbf{x}) = \sum_{i=1}^{n_{point}} \sum_{j=1}^{n_{var}} \left[ y_{i,j}^{sim}(\mathbf{x}) - y_{i,j}^{exp} \right]^2$, where **x** is the model parameter vector (the decision variables in the terminology of optimization problems). $y_{i,j}^{sim}$ and $y_{i,j}^{exp}$ indicate the simulated and experimental data, respectively. $n_{point}$ and $n_{var}$ are the number of data points and the number of model variables, respectively. Note that by default, the objective function for parameter estimation supports only the *unconstrained* optimization formulation. That is, the objective function returns the squared sum of the differences between simulated data and experimental data as $f$ and does not return $g_i$. The functions **RCGA_UNDXMGG_PE** and **RCGA_REXstarJGG_PE** return the structure **Results** in the same way that **RCGA_UNDXMGG** and **RCGA_REXstarJGG** does.

After starting the RCGAs, a pop-up figure appears which will update during the calculation (**Figure 1**). In the figure, circles and lines indicate the target experimental data and the current best model behavior, respectively. The model fitting improves as the calculations progress.

### 4. Tutorial

#### 4.1. Solving general optimization problems

##### 4.1.1. Execution via script files

Having explained the basic usage of RCGAToolbox, we explain its advanced usage in this section. RCGAToolbox/doc/demo/general/**run_Tutorial.m** is the script to solve the same constrained optimization problem as that in **Section 3.1** but with different options.

RCGAToolbox/doc/demo/general/run_Tutorial.m

```
% This script demonstrates how to run a real-coded genetic algorithm to
% solve an example constrained optimization problem.
%
% --------------------- Example Problem ---------------------
% Minimize:
%   f = x(1)^2 + x(2)^2 + ... + x(10)^2
%
% Subject to:
%   g(1) = x(1) * x(2) + 1 <= 0
%   g(2) = x(1) + x(2) + 1 <= 0
%   -5.12 <= x(i) <= 5.12 for all i
%
%
```

```
% Global minimum is f = 3, g(1) = 0, g(2) = 0 at x = (-1.618, 0.6180, 0, 0,
% 0, 0, 0, 0, 0, 0) or at x = (0.6180, -1.618, 0, 0, 0, 0, 0, 0, 0, 0)
% -----------------------------------------------------------

clearvars;

% ========= Problem Settings ========= %
problem.n_gene = 10; % Number of Decision Variables
problem.n_constraint = 2; % Number of Constraints
problem.fitnessfun = @Fitness_Example; % Fitness Function
problem.decodingfun = @Decoding_Example; % Decoding Function

% ========= Option Settings ========== %
opts.n_population = 100; % Population Size
opts.n_children = 100; % Number of Children per Generation
opts.n_parent = problem.n_gene + 1; % Number of Parents Used for REXstar
opts.t_rexstar = 6.0; % Step-size Parameter for REXstar
opts.selection_type = 0; % Parameter for JGG (0: Chosen from Children, 1: Chosen
from Family)
opts.Pf = 0.45; % Pf
opts.local = 0; % Local Optimizer
% opts.localopts =
optimoptions(@fmincon,'ConstraintTolerance',0,'MaxFunctionEvaluations',opts.n_ch
ildren,'Display','off'); % Options for Local Optimizer
opts.maxgen = 1000; % Max Number of Generations
opts.maxtime = 60; % Max Time (sec)
opts.maxeval = inf; % Max Number of fitnessfun Evaluations
opts.vtr = 0; % Value To Be Reached
opts.n_par = 1; % Number of Workers for Parallel Computation
opts.output_intvl = 10; % Output Interval Generation
opts.out_transition = 'Transition.txt'; % Transition File Name
opts.out_best = 'BestIndividual.txt'; % Best Individual File Name
opts.out_population = 'FinalPopulation.txt'; % Final Population File Name
opts.out_report = 'Report.mat'; % Report File Name
opts.interimreportfun = @RCGAdefaultinterimreportfun; % Interim Report Function
opts.finalreportfun = @RCGAdefaultfinalreportfun; % Final Report Function
```

```
% ======= Setting Random Seed ======== %
rng(0); % Random Seed

% ========== Executing RCGA ========== %
% Results = RCGA_UNDXMGG(problem,opts); % UNDX/MGG
Results = RCGA_REXstarJGG(problem,opts); % REXstar/JGG
```

As shown in **run_Tutorial.m**, the following fields of the structure **opts** can be set:

- **opts.n_population**: Population size, that is, the number of individuals in the population. (Default: 300)
- **opts.n_children**: Number of children that are generated per generation. (Default: **opts.n_population**)
- **opts.n_parent**: Number of parents that are used to generate the children. This option is active only for $REX^{star}$/JGG. (default: **problem.n_gene** + 1)
- **opts.t_rexstar**: Step size parameter. This option is only active for $REX^{star}$/JGG. (Default: 6.0)
- **opts.selection_type**: Selection type. **opts.selection_type** must be 0 or 1. If it is 0, a certain number (**opts.n_parent**) of the best children are selected. If it is 1, a certain number (**opts.n_parent**) of individuals in the family (children and parents) are selected. This option is only active for $REX^{star}$/JGG. (Default: 0)
- **opts.Pf**: Probability that only the objective function $f$ is used for comparisons of individuals in the stochastic ranking. This option is only active for constrained optimization problems (**problem.n_constraint** > 0). The recommended value is $0.4 < P_f < 0.5$. (Default: 0.45)
- **opts.local**: Local optimizer. **opts.local** must be 0 or 1. If it is 1, the local optimizer is used: when the best individual is updated after generation alternation by UNDX/MGG or $REX^{star}$/JGG, the local optimizer is called to further improve the best individual. Whether the use of the local optimizer improves the performance of RCGAs depends on the specific problems. Generally, in problems with multiple local optima, the local optimizer does not work well, and thus, its use leads to a higher computational cost. Note that Optimization Toolbox is required for the local optimizer. (Default: 0)
- **opts.localopts**: Options for the local optimizer. The options are active only when **opts.local** = 1. (Default: optimoptions( @fmincon, ’ConstraintTolerance’, 0, 'MaxFunctionEvaluations', opts.n_children, 'Display', 'off' ); Moreover, “UseParallel” is set to “true” if **opts.n_par** > 1)
- **opts.maxgen**: Number of maximum generations. RCGAs end once they reach **opts.maxgen**. (Default: 1000)

- **opts.maxtime**: Maximum time (sec). RCGAs end when they reach **opts.maxtime**. (Default: 600)
- **opts.maxeval**: Maximum number of fitness function evaluations. RCGAs end when they reach **opts.maxeval**. (Default: Inf)
- **opts.vtr**: Value to be reached. RCGAs end when they reach **opts.vtr**. (Default: –Inf)
- **opts.n_par**: Number of workers in parallel computation. In parallel computation, each worker evaluates **opts.n_population/opts.n_par** individuals in the first generation and **opts.n_children/opts. n_par** individuals in the second generation to the final generation. Thus, the more workers, the fewer individuals to be evaluated per worker, leading to faster progression of RCGAs. Note that Parallel Computing Toolbox is required for parallel computation (**opts.n_par** > 1). (Default: 1)
- **opts.output_intvl**: The number of generations after which the transition file and the report file are each time updated. When this option is properly applied, users can avoid generating huge output files. (Default: 1)
- **opts.out_transition**: Name of an output file called the transition file. The transition file is a text file in which each row corresponds to the best individual in each generation. The columns show the computational time (*Time*), the number of fitness function evaluations (*NEval*), generation (*Generation*), objective function value (*f*), penalty function (*phi*), decision variables (*x*), and constraint function values (*g*). (Default: 'None')
- **opts.out_best**: Name of an output file called the best individual file. The best individual file is a text file that is generated at the end of RCGAs and contains the information on the best individual found in the computation. It also contains the same columns as the transition file. (Default: 'None')
- **opts.out_population**: Name of an output file called the final population file. The final population file is a text file that is generated at the end of RCGAs and contains information on all individuals in the final population. Each row corresponds to each individual. The columns are the objective function value (*f*), penalty function (*phi*), decision variables (*x*), and constraint function values (*g*). (Default: 'None')
- **opts.out_report**: Name of an output file called the report file. The report file is a MATLAB MAT file in which the structure **Results** is stored. The structure **Results** is the same as the structure **Results** returned by **RCGA_UNDXMGG** and **RCGA_REXstarJGG** at the end of the RCGAs. The structure contains four fields: **Transition**, **Best**, **FinalPopulation**, and **end_crit**. The field **Transition** is updated at each **opts.interval** generation, while the other fields are generated at the end of the RCGAs. (Default: 'None')
- **opts.interimreportfun**: Function handle of the interim report function called at each **opts.output_intvl** generation. The function **RCGAdefaultinterimreportfun** can be used as a template for custom interim report functions. (Default: @RCGAdefaultinterimreportfun)
- **opts.finalreportfun**: Function handle of the final report function called at the end of RCGAs. The

function **RCGAdefaultfinalreportfun** can be used as a template for custom final report functions. (Default: @RCGAdefaultfinalreportfun)

In **run_Tutorial.m**, the function **rng** is used for reproducibility. If **rng** is not called before starting RCGAs, the results will change each time users run RCGAs. Executing **run_Tutorial.m** provides similar messages to **run_QuickStart.m** in the MATLAB Command Window. Four output files (Transition.txt, BestIndividual.txt, FinalPopulation.txt, and Report.mat) are generated in the directory RCGAToolbox/doc/demo/general/.

###### 4.1.2. Execution via GUI

For those unfamiliar with codes or scripts, the RCGAToolbox has graphical user interfaces (GUIs). To start the GUI "RCGAToolbox Mission Control", execute **run_GUI.m** under the directory RCGAToolbox/doc/demo/general/, or alternatively type "RCGAToolbox_MissionControl" in the MATLAB Command Window. Using the GUI, users can specify problems and options (**Figure 2**). To run RCGAs, click the "Launch" button.

Most of the items in the GUI correspond to the structures **problem** and **opts** explained above. The "Save" button saves the current settings in a MAT file. The "Load" button allows loading the settings from a created MAT file. By clicking the "Reset" button, all the settings in the GUI can be reset. If you are not familiar with RCGAs, you can use recommended settings by clicking the "Use Recommended Values" button. By clicking the "Create" button, the GUI creates an executable script (a file similar to **run_Tutorial.m**) with the current settings. We recommend that beginners first use the GUI. After getting comfortable with RCGAs, they can create the executable script by the "Create" button and modify it for advanced use.

##### 4.2. Solving parameter estimation problems in systems biology

Examples in this section require IQM Tools. If you have not installed IQM Tools, please use sample scripts in RCGAToolbox/doc/demo/*PE_woIQM*/ instead of RCGAToolbox/doc/demo/*PE*/.

###### 4.2.1. Execution via script files

RCGAToolbox/doc/demo/PE/**run_Tutorial.m** is another script to solve the parameter estimation problem found in **Section 3.2** but with different options.

RCGAToolbox/doc/demo/PE/run_Tutorial.m

```
% This script demonstrates how to run a real-coded genetic algorithm to
% estimate model parameters in an example kinetic model.
%
% ------------------------ Example Kinetic Model ------------------------
% - INITIAL CONDITION
% X1 = 0
% X2 = 0
%
% - PARAMETERS
% X0 = 0.1
% k1 = 1
% k2 = 1
% k3 = 1
% K2 = 1
% K3 = 1
% rootCompartment = 1
%
% - VARIABLES
% X12 = X1 + X2
%
% - REACTIONS
% v1 = k1 * X0
% v2 = k2 * (X1/rootCompartment) / (K2 + (X1/rootCompartment))
% v3 = k3 * (X2/rootCompartment) / (K3 + (X2/rootCompartment))
%
% - BALANCE
% X1_dot = v1 - v2;
% X2_dot = v2 - v3;
% -----------------------------------------------------------------------

clear mex;
clear all;
close all;

% ========= Problem Settings ========= %
```

```
% modelfile = 'Model_Example_SBML.xml'; % SBML file (IQM Tools required)
% modelfile = IQMmodel('Model_Example_SBML.xml'); % Creating an IQMmodel object
(IQM Tools required)
modelfile = @Model_Example_odefun; % ODE file (IQM Tools format)
% modelfile = 'Model_Example_odefun.m'; % ODE file (IQM Tools format)
% modelfile = 'Model_Example_mex.c'; % C source code (IQM Tools required)
% modelfile = 'Model_Example_mex.mexw64'; % MEX model for Windows
% modelfile = 'Model_Example_mex.mexmaci64'; % MEX model file for macOS
% modelfile = 'Model_Example_mex.mexa64'; % MEX model file for Linux
decodingfun = @Decoding_Example; % Decoding Function
% measurement = 'Measurement_Example.xls'; % Measurement File (EXCEL format)
measurement = 'Measurement_Example.csv'; % Measurement File (CSV)

% ========= Option Settings ========== %
opts.n_population = 50; % Population Size
opts.n_children = 25; % Number of Children per Generation
opts.n_parent = 7 + 1; % Number of Parents Used for REXstar
opts.t_rexstar = 6.0; % Step-size Parameter for REXstar
opts.selection_type = 0; % Parameter for JGG (0: Chosen from Children, 1: Chosen
from Family)
opts.local = 0; % Local Optimizer
% opts.localopts =
optimoptions(@fmincon,'ConstraintTolerance',0,'MaxFunctionEvaluations',opts.n_ch
ildren,'Display','off'); % Options for Local Optimizer
opts.maxgen = 200; % Max Number of Generations
opts.maxtime = 60; % Max Time (sec)
opts.maxeval = inf; % Max Number of fitnessfun Evaluations
opts.vtr = 0; % Value To Be Reached
opts.n_par = 1; % Number of Workers for Parallel Computation
opts.output_intvl = 1; % Output Interval Generation
opts.out_transition = 'Transition.txt'; % Transition File Name
opts.out_best = 'BestIndividual.txt'; % Best Individual File Name
opts.out_population = 'FinalPopulation.txt'; % Final Population File Name
opts.out_report = 'Report.mat'; % Report File Name
fast_flag = 0; % fast_flag (0: MATLAB ODEXX)
% fast_flag = 1; % fast_flag (1: SundialsTB CVODE) (SundialsTB required)
% fast_flag = 2; % fast_flag (2: IQM Tools CVODE MEX) (IQM Tools required)
```

```
% ======= Setting Random Seed ======== %
rng(0); % Random Seed

% ========== Executing RCGA ========== %
% Results =
RCGA_UNDXMGG_PE(modelfile,decodingfun,measurement,fast_flag,[],opts); % UNDX/MGG
Results =
RCGA_REXstarJGG_PE(modelfile,decodingfun,measurement,fast_flag,[],opts); %
REXstar/JGG
```

The following three are the inputs specific for parameter estimation:

- **modelfile**: Name of a model file or IQMmodel object. The model file can be an SBML file, an ODE file (IQM Tools format), a C source code, or a MEX model. The IQMmodel object is a MATLAB object used in IQM Tools. (For details, see the user guide of IQM Tools).
- **measurement**: Measurement file that provides experimental data to be fitted by the model. The measurement file can be a CSV file or an XLS file (for details, see the user guide of IQM Tools).
- **fast_flag**: Input specifying which ODE solver is used. For **fast_flag** = 0, a built-in MATLAB solver is used. For **fast_flag** = 1, a pre-compiled ODE solver (CVODE) provided by SundialsTB is used. For **fast_flag** = 2, a pre-compiled ODE solver (CVODE) provided by IQM Tools is used with a MEX model.

##### 4.2.2. Execution via GUI

RCGAToolbox has a GUI for parameter estimation (**Figure 3**). To start the GUI "RCGAToolbox Mission Control PE", execute **run_GUI.m** in the directory RCGAToolbox/doc/demo/PE/, or type "RCGAToolbox_MissionControl_PE" in the MATLAB Command Window. Most input fields shown in the GUI are the same as those in the GUI for general use (see **Section 4.1.2**). Additional input fields are **modelfile**, **measurement**, and **fast_flag**.

#### 4.3. Simulating kinetic models

RCGAToolbox/doc/demo/sim/**run_SimulateModel.m** demonstrates how to simulate a kinetic model. A kinetic model can be provided as an SBML file, an IQMmodel object, an ODE file (IQM Tools format), a C source code, or a MEX model (In **run_SimulateModel.m**, you can comment/uncomment the lines starting with "model ="). An IQMmodel object is an object that represents a kinetic model in IQM Tools (for details, see the user guide of IQM Tools). The function **RCGAsimulate** simulates the model with an ODE solver specified by **fast_flag** (you can comment/uncomment the lines starting with "fast_flag ="). When **fast_flag** = 0, MATLAB built-in ODE solvers are used. When **fast_flag** = 1, a pre-compiled ODE solver provided by SundialsTB is used. When **fast_flag** = 2, the model is compiled by IQM Tools and combined with a pre-compiled ODE solver. As **run_SimulateModel.m** is executed, a figure pops up, which shows the model behavior. The function **RCGAsimulate** is useful because it checks the model format and **fast_flag**, and then automatically chooses an appropriate ODE solver. For **fast_flag** = 1 and 2, SundialsTB and IQM Tools are required, respectively. For the function IQMmodel, IQM Tools is required.

RCGAToolbox/doc/demo/sim/run_SimulateModel.m

```
% This script demonstrates how to run simulation using RCGAToolbox.

clear mex;
clear all;
close all;

% =============== Model =============== %
% model = 'Model_Example_SBML.xml'; % SBML file (IQM Tools required)
% model = IQMmodel('Model_Example_SBML.xml'); % Creating an IQMmodel object (IQM
Tools required)
model = @Model_Example_odefun; % ODE file (IQM Tools format)
% model = 'Model_Example_odefun.m'; % ODE file (IQM Tools format)
% model = 'Model_Example_mex.c'; % C source code (IQM Tools required)
% model = 'Model_Example_mex.mexw64'; % MEX model for Windows
% model = 'Model_Example_mex.mexmaci64'; % MEX model file for macOS
% model = 'Model_Example_mex.mexa64'; % MEX model file for Linux

% =============== Time ================ %
tspan = 0 : 0.1 : 10;

% ========= Initial Condition ========= %
y0(1) = 0; % X1
y0(2) = 0; % X2

% ========= Parameter Values ========== %
param(1) = 0.1; % X0
```

```
param(2) = 1;   % k1
param(3) = 1;   % k2
param(4) = 1;   % k3
param(5) = 1;   % K2
param(6) = 1;   % K3
param(7) = 1;   % rootCompartment

% ============ ODE Solver ============= %
fast_flag = 0; % # fast_flag (0: MATLAB ODEXX)
% fast_flag = 1; % # fast_flag (1: SundialsTB CVODE) (SundialsTB required)
% fast_flag = 2; % # fast_flag (2: IQM Tools CVODE MEX) (IQM Tools required)

% ============ Simulation ============= %
[ T, Y ] = RCGAsimulate(model, tspan, y0, param, fast_flag);

% =============== Figure ============== %
figure;
plot(T,Y,'-','LineWidth',2);
legend('X_1','X_2','Location','best');
xlabel('Time');
ylabel('Concentration');
```

### 5. Useful features

#### 5.1. Model format conversion

RCGAToolbox/doc/demo/conversion/**run_Conversion.m** demonstrates how to convert an IQMmodel object, an SBML file, an ODE file, a concise ODE file, a C source code, and a MEX model into one another. RCGAToolbox, combined with IQM Tools, provides different conversion functions, as summarized in **Figure 4**. The script **run_Conversion.m** creates different model files from the original model file **Model_Example_conciseOdefun.m**, which is written in the concise ODE file format as shown below.

RCGAToolbox/doc/demo/conversion/Model_Example_conciseOdefun.m

```
function dydt = Model_Example_conciseOdefun(t, y)
% Model_Example_conciseOdefun is an example of the concise ODE file
% (RCGAToolbox format) ready for conversion into an IQMmodel object by
% RCGAreadConciseODEfile. In this file, the only sections sandwiched
% between "BEGIN" and "END" are used for the conversion.
%
% [SYNTAX]
% dydt = Model_Example(t, y)
%
% [INPUT]
% t    :  Time.
% y    :  X1 and X2.
%
% [OUTPUT]
% dydt :  dX1/dt and dX2/dt.

X1 = y(1);
X2 = y(2);

% === BEGIN NAME ===
% Model_Example
% === END NAME ===

% === BEGIN NOTES ===
% Simple metabolic pathway with two Michaelis-Menten rate equations.
% === END NOTES ===

% === BEGIN INITIAL CONDITION ===
X1_0 = 0;
X2_0 = 0;
% === END INITIAL CONDITION ===

% === BEGIN PARAMETERS ===
X0 = 0.1;
k1 = 1;
```

```
k2 = 1;
k3 = 1;
K2 = 1;
K3 = 1;
% === END PARAMETERS ===

% === BEGIN VARIABLES ===
X12 = X1 + X2;
% === END VARIABLES ===

% === BEGIN REACTIONS ===
v1 = k1 * X0;
v2 = k2 * X1 / ( K2 + X1 );
v3 = k3 * X2 / ( K3 + X2 );
% === END REACTIONS ===

% === BEGIN BALANCE ===
X1_dot = v1 - v2;
X2_dot = v2 - v3;
% === END BALANCE ===

dydt = zeros(2,1);
dydt(1) = X1_dot;
dydt(2) = X2_dot;
```

The concise ODE file format is simpler than the ODE file format used in IQM Tools (RCGAToolbox/doc/demo/sim/**Model_Example_odefun.m** is written in the IQM Tools ODE file format). The concise ODE file can be used for ODE solvers and yet ready for conversion to SBML. Users can easily implement their models in the concise ODE file format, and once done, those models can be automatically converted into IQMmodel objects or SBML files. IQM Tools are required for file conversion functions.

##### 5.2. Wrapper for the fast ODE solver CVODE

RCGAToolbox/doc/demo/stb/**run_ODESTB.m** demonstrates how to use the fast ODE solver CVODE. In general, the ODE solver CVODE provided by SundialsTB is faster than MATLAB's built-in ODE solvers such as ode15s and ode23s (see **Section 7.1**). However, the use of CVODE is more complicated than that of MATLAB's built-in ODE solvers. The function **odestb** provided by RCGAToolbox is a wrapper for CVODE and enables the use of CVODE in the same way as the built-in ODE solvers are used. SundialsTB is required to run **run_ODESTB.m**.

RCGAToolbox/doc/demo/stb/run_ODESTB.m

```
% This script demonstrates how to use CVODE by SundialsTB. SundialsTB is
% required to use the function odestb.

clear mex;
clear all;
close all;

% =============== Time ================ %
tspan = 0 : 0.1 : 10;

% ========= Initial Condition ========= %
y0(1) = 0; % X1
y0(2) = 0; % X2

% ============ Simulation ============= %
tic;
% [ T, Y ] = ode45(@Model_Example_conciseOdefun, tspan, y0); % MATLAB Built-in
% [ T, Y ] = ode23(@Model_Example_conciseOdefun, tspan, y0); % MATLAB Built-in
% [ T, Y ] = ode113(@Model_Example_conciseOdefun, tspan, y0); % MATLAB Built-in
% [ T, Y ] = ode15s(@Model_Example_conciseOdefun, tspan, y0); % MATLAB Built-in
% [ T, Y ] = ode23s(@Model_Example_conciseOdefun, tspan, y0); % MATLAB Built-in
% [ T, Y ] = ode23t(@Model_Example_conciseOdefun, tspan, y0); % MATLAB Built-in
% [ T, Y ] = ode23tb(@Model_Example_conciseOdefun, tspan, y0); % MATLAB Built-in
[ T, Y ] = odestb(@Model_Example_conciseOdefun, tspan, y0); % CVODE provided by
SundialsTB
toc

% =============== Figure ============== %
figure;
plot(T,Y,'-','LineWidth',2);
```

```
legend('X_1','X_2','Location','best');
xlabel('Time');
ylabel('Concentration');
```

#### 5.3. Implementing custom RCGAs

RCGAToolbox enables users to efficiently implement their custom RCGAs [or other evolution computation algorithms such as the evolution strategy (Runarsson and Yao, 2005), differential evolution (Takahama and Sakai, 2010), and scatter search (Egea, et al., 2009)]. Users need to provide a user-defined generation alternation function that defines how to reproduce new individuals (children) and how to update the population. The script **run_CustomRCGA.m** demonstrates how to run user-defined custom RCGAs.

RCGAToolbox/doc/demo/CustomRCGA/run_CustomRCGA.m

```
% This script demonstrates how to run a user-defined custom real-coded
% genetic algorithm to solve an example constrained optimization problem.
%
% --------------------- Example Problem ---------------------
% Minimize:
%   f = x(1)^2 + x(2)^2 + ... + x(10)^2
%
% Subject to:
%   g(1) = x(1) * x(2) + 1 <= 0
%   g(2) = x(1) + x(2) + 1 <= 0
%   -5.12 <= x(i) <= 5.12 for all i
%
%
% Global minimum is f = 3, g(1) = 0, g(2) = 0 at x = (-1.618, 0.6180, 0, 0,
% 0, 0, 0, 0, 0, 0) or at x = (0.6180, -1.618, 0, 0, 0, 0, 0, 0, 0, 0)
% -----------------------------------------------------------

clearvars;

% ========= Problem Settings ========= %
problem.n_gene = 10; % Number of Decision Variables
problem.n_constraint = 2; % Number of Constraints
```

```
problem.fitnessfun = @Fitness_Example; % Fitness Function
problem.decodingfun = @Decoding_Example; % Decoding Function

% ========== Executing RCGA ========== %
% GenerationAlternation_Example is a user-defined custom generation
% alternation function
Results = RCGA_CustomRCGA(problem, @GenerationAlternation_Example);
```

As you can see, the function **RCGA_CustomRCGA** receives the structure **problem** and the function handle for a generation alternation function. If necessary, users can provide **RCGA_CustomRCGA** with the structure **opts**. **GenerationAlternation_Example.m** is a sample generation alternation function and can be used as a template for custom generation alternation functions.

RCGAToolbox/doc/demo/CustomRCGA/GenerationAlternation_Example.m

```
function Population = GenerationAlternation_Example(problem, opts, Population)
% GenerationAlternation_Example updates population by a simple generation
% alternation algorithm. This function can be used as a template for custom
% generation alternation functions.
% ...

%% Shortening variable names
n_population = opts.n_population;
n_children = n_population - 1; % Note that opts.n_children is not used.
n_constraint = problem.n_constraint;
n_gene = problem.n_gene;
Pf = opts.Pf;

%% Generating new children
ip = randperm(n_population,2);

c(1,1:n_children) =
struct('gene',zeros(1,n_gene),'g',zeros(1,n_constraint),'f',zeros,'phi',zeros);

for i = 1 : n_children
```

```
    % Crossover
    c(i).gene = 0.5 * ( Population(ip(1)).gene + Population(ip(2)).gene );

    % Mutation
    mutation_rate = 0.1;
    for j = 1 : n_gene
        if rand < mutation_rate
            c(i).gene(j) = rand;
        end
    end

end
c = RCGArequestFitnessCalc(problem,opts,c);

%% Making a new population
Population = [ Population(1) c ];
Population = RCGAsrsort(Population,Pf);
```

The input arguments are the structure **problem**, structure **opts**, and structure array **Population**. An element of **Population** is the structure **individual** that has the vector **gene**, vector **g**, scaler **f**, and scaler **phi** as fields. The function **GenerationAlternation_Example** produces new individuals (children) based on the given population. Primitive crossover and mutation methods are used for the sake of simplicity. **GenerationAlternation_Example** keeps the best individual in the population and replaces the other individuals with the children. The predefined function **RCGArequestFitnessCalc** calculates values of the objective function (*f*), constraint functions (*g*), and penalty function (*phi*) for the children. The predefined function **RCGAsrsort** sorts individuals in the population according to the stochastic ranking sort algorithm.

### 6. Real-coded genetic algorithms

Genetic algorithms (GAs) are metaheuristic techniques developed with inspiration from the evolution of living organisms (Reali, et al., 2017; Sun, et al., 2012). The basic procedure is as follows:

1. Generate an initial population in which each individual is characterized by a set of different values for the decision variables.
2. Select a subset of individuals (called parents) from the population.
3. Generate children using the selected parents (outside the population).
4. Select children that show good fitness (i.e., featuring small values of the objective function).
5. Replace the parents in the population with the same number of selected children, thereby creating a partly changed population, while maintaining the total number of individuals.
6. If a termination criterion is not met by the new population, return to step 2.

The above procedure is terminated when a sufficiently good individual is obtained or the number of generations reaches the predefined maximal number. Various GAs have been proposed so far, and they differ in their implementation for each step. For parameter estimation in systems biology, real-coded GAs (RCGAs) are used, in which a set of kinetic parameters (i.e., a real number vector) is handled as an individual in steps 1-6. In parameter estimation, a good fitness in step 4 means good fitting to the experimental data for training the model.

In constrained optimization problems, it is essential to balance the objective function and the constraint functions for the appropriate ranking of individuals. To achieve a well-balanced ranking, the stochastic ranking method based on the bubble-sort-like procedure was proposed (**Figure 5**). For details on constraint handling, see section 2.3.3 of our previous study (Maeda, et al., 2018).

#### 6.1. UNDX/MGG

UNDX/MGG employs unimodal normal distribution crossover (UNDX) (Ono and Kobayashi, 1997) as the crossover method and minimal generation gap (MGG) (Satoh, et al., 1997) as the generation alternation method. The UNDX and MGG algorithms are described below. Here, $n_v$ is the number of decision variables, and $n_c$ is the number of children. In the UNDX, there are two parameters: $\alpha$ and $\beta$. We followed the original recommendations (Ono and Kobayashi, 1997) of $\alpha$ = 0.5 and $\beta$ = 0.35. It should be noted that UNDX uses three parents, but MGG was originally developed for two-parent-based crossover. To combine MGG and UNDX, the subparent ($\mathbf{x}_3$) in UNDX is randomly chosen from the population but not treated as a parent in MGG.

The algorithm of MGG is described as follows:

1. Randomly select two individuals from the population.
2. Using these two individuals (hereafter "parents"), create $n_c$ children.
3. Calculate the objective function $f$ and the penalty function $\phi$ for the children.
4. Replace the two parents in the population with the best individual and a randomly-chosen individual from the "family." The family consists of parents and children.

The algorithm of UNDX is described as follows:

1. Two main parents $\mathbf{x}_1$ and $\mathbf{x}_2$ and a single sub parent $\mathbf{x}_3$ are used for the crossover.
2. Calculate the midpoint of $\mathbf{x}_1$ and $\mathbf{x}_2$: $\mathbf{x}_p = (\mathbf{x}_1 + \mathbf{x}_2)/2$.
3. Calculate the vector from $\mathbf{x}_1$ to $\mathbf{x}_2$: $\mathbf{d} = \mathbf{x}_2 - \mathbf{x}_1$.
4. Calculate the distance ($D$) between $\mathbf{x}_3$ and the primary search line (the line passing through $\mathbf{x}_1$ and $\mathbf{x}_2$).
5. Generate a child $\mathbf{x}_c$ using the equation below:

$$\mathbf{x}_c = \mathbf{x}_p + \xi\mathbf{d} + D\sum_{i=1}^{n_v-1}\eta_i\mathbf{e}_i\,, \qquad (3)$$

where $\xi \sim N\left(0, \sigma_\xi^2\right)$, $\eta_i \sim N\left(0, \sigma_\eta^2\right)$, and $\mathbf{e}_i$ is the $i^{\text{th}}$ orthogonal basis vector. $\sigma_\xi = \alpha$ and $\sigma_\eta = \beta / \sqrt{n_v}$. Multiple children can be generated by applying Eq. (3) multiple times. $N(x, y)$ is the normal random distribution with the mean $x$ and standard deviation $y$.

#### 6.2. REX$^{\text{star}}$/JGG

REX$^{\text{star}}$/JGG employs real-coded ensemble crossover star (REX$^{\text{star}}$) as the crossover method and just generation gap (JGG) as the generation alternation method (Kobayashi, 2009). The algorithms of REX$^{\text{star}}$ and JGG are described below. Here, $n_v$ is the number of decision variables, $n_p$ is the number of parents, $n_c$ is the number of children, and $t$ is the step-size parameter. $t$ determines the aggressiveness of the search. $n_p \leq n_c$ must be satisfied to maintain a constant population size. In the original study, $n_p = n_v + 1$ is recommended, and $t$ is set between 2.5 and 15 (Kobayashi, 2009).

The algorithm of JGG is described as follows:

1. Randomly select $n_p$ individuals from the population.
2. Using these selected individuals (hereafter "parents"), create $n_c$ children.
3. Calculate the objective function $f$ and the penalty function $\phi$ for the children.
4. Replace the parents in the population with the best $n_p$ children.

The algorithm of REX$^{star}$ is described as follows:

1. $n_p$ parents $\mathbf{x}_1$, $\mathbf{x}_2$, ..., $\mathbf{x}_{n_p}$ are used for the crossover.
2. Generate the reflection points $\mathbf{x}'_1$, $\mathbf{x}'_2$, ..., $\mathbf{x}'_{n_p}$ of the parents using Eqs. (4) with the center of gravity $\mathbf{x}_g$:

$$\mathbf{x}_g = \frac{1}{n_p}\sum_{i=1}^{n_p}\mathbf{x}_i \text{ ,} \quad (4a)$$

$$\mathbf{x}'_i = 2\mathbf{x}_g - \mathbf{x}_i \text{ .} \quad (4b)$$

3. Calculate the objective function $f$ and the penalty function $\phi$ for the reflection points.
4. Select the best $n_p$ individuals from the parents and their reflection points.
5. Compute the center of gravity ($\mathbf{x}_b$) of the selected individuals.
6. Generate a child $\mathbf{x}_c$ using the equation below:

$$\mathbf{x}_c = \mathbf{x}_g + \mathrm{diag}(\xi_1^t, \cdots, \xi_{n_v}^t)\cdot(\mathbf{x}_b - \mathbf{x}_g) + \sum_{i=1}^{n_p}\xi^i(\mathbf{x}_i - \mathbf{x}_g)\text{ ,} \quad (5)$$

where $\xi_i^t \sim U(0,t)$ ($i$ = 1, ..., $n_v$), and $\xi^i \sim U\left(-\sqrt{3/n_p}, \sqrt{3/n_p}\right)$ ($i$ = 1, ..., $n_p$). Multiple children can be generated by applying Eq. (5) multiple times. $U(x, y)$ is the uniform random distribution with the lower bound $x$ and the upper bound $y$.

### 7. Computational experiments

#### 7.1. Comparison of simulation methods

In parameter estimation, kinetic models are simulated every time the objective function and the constraint functions are calculated. Thus, it is essential to accelerate simulations for rapid parameter estimation. The RCGAToolbox provides access to two fast third-party solvers (CVODE provided by SundialsTB and IQM Tools).

We performed a computational experiment to investigate how much RCGAToolbox accelerates simulations by fast solvers. The results are summarized in **Figure 6**. The MATLAB built-in solver (**fast_flag** = 0) is the slowest, and CVODE provided by IQM Tools (**fast_flag** = 2) is the fastest. The CVODE provided by SundialsTB (**fast_flag** = 1) is in between. Depending on the model, CVODE by IQM Tools (**fast_flag** = 2) is more than 30 times faster than MATLAB's built-in solver (**fast_flag** = 0). Therefore, we recommend using CVODE by IQM Tools (**fast_flag** = 2) if available. In the directory RCGAToolbox/doc/benchmark/sim, the scripts for the computational experiment are provided, which enable interested readers to perform computational experiments on their computers. Please note that the results can vary depending on the computational environment.

##### 7.2. Unconstrained test problems

To evaluate the performance of the RCGAToolbox in unconstrained optimization problems, we employed 14 test problems (**Appendix C**). Except for Sphere, these problems have one or more properties that hamper the search (i.e., ill-scale, intervariable dependency, and multimodality). Sphere does not have such properties and thus can be considered a control. We define the solutions as decision variable vectors that provide $f \leq 10^{-6}$.

We applied the UNDX/MGG and $\text{REX}^{\text{star}}$/JGG provided by RCGAToolbox to the 14 unconstrained test problems. As a control, we used MATLAB GA provided by Global Optimization Toolbox (MathWorks, Inc., Natich, MA, USA). As another control, we employed eSS (Enhanced Scatter Search) (Egea, et al., 2009; Pardo, et al., 2020). Specifically, we used eSS (R2019) provided in the latest version of MEIGO (Egea, et al., 2014). We used fmincon as the local solver for the UNDX/MGG, $\text{REX}^{\text{star}}$/JGG, and eSS. We ran each optimization algorithm 10 times per problem. We set the maximum computational time to 30 min for each run.

The number of successful trials is summarized in **Table 1**. MATLAB GA found solutions only in 3 out of 14 problems. UNDX/MGG from RCGAToolbox found solutions in four problems. Both eSS and $\text{REX}^{\text{star}}$/JGG found solutions in 12 problems. Rosenbrock (star) and Schwefel were solved only by eSS. Schaffer and the scaled shifted rotated Rastrigin were solved only by $\text{REX}^{\text{star}}$/JGG. In summary, $\text{REX}^{\text{star}}$/JGG and eSS worked comparatively well and outperformed MATLAB GA and UNDX/MGG. In the directory RCGAToolbox/doc/benchmark/math, the scripts for the computational experiments are provided. Please note that all algorithms were used with default settings (except termination criteria and local solvers). The results can vary when using different settings.

##### 7.3. Constrained test problems

To evaluate the performance of the RCGAToolbox in constrained optimization problems, we employed 13 test problems (**Appendix D**). For the constrained test problems, we define solutions as being decision variable vectors that provide an $f$ value close to the known optimal solution with $\phi = 0$. We set the allowable error for $f$ to 1%.

We applied the UNDX/MGG and $\text{REX}^{\text{star}}$/JGG provided by RCGAToolbox to the 13 constrained test problems. As controls, we employed MATLAB GA and eSS, as described in **Section 7.2**. We used fmincon as the local solver for the UNDX/MGG, $\text{REX}^{\text{star}}$/JGG, and eSS. We ran each optimization algorithm 10 times per problem. We set the maximum computational time to 30 min for each run.

The number of successful trials is summarized in **Table 2**. MATLAB GA found solutions for 5 out of 13 problems. UNDX/MGG, $REX^{star}$/JGG, and eSS worked comparatively well and found solutions for all 13 problems. In the directory RCGAToolbox/doc/benchmark/math, the scripts for the computational experiment are provided. Please note that the results can vary depending on the options for the optimization algorithms.

#### 7.4. Biological test problems

To evaluate the performance of RCGAToolbox in parameter estimation in systems biology, we employed six test problems (**Table 3**): two from our previous study (Maeda, et al., 2018) and four from the BioPreDyn-Bench benchmark suit (Villaverde, et al., 2015). These test problems are parameter estimation problems for medium to large-scale kinetic models (20 - 117 unknown parameters). The objective function is defined as the (weighted) difference between model prediction and experimental data. The three-step and HIV problems are constrained optimization problems (see **Appendix E**), whereas B2, B4, B5, and B6 are unconstrained optimization problems (i.e., no constraint functions). We did not employ B1 and B3 of BioPreDyn-Bench since B1 is computationally expensive and B3 did not run on our environment. We employed values to be reached (VTR, the objective function value below which decision variable vectors are regarded as solutions) as indicated in **Table 3**, so that models with solutions provide a sufficiently good fit to experimental data.

We applied UNDX/MGG and $REX^{star}$/JGG provided by RCGAToolbox to the six parameter estimation problems. We employed MATLAB GA and eSS (R2019) as controls. We used fmincon as the local solver for UNDX/MGG, $REX^{star}$/JGG, and eSS. We ran each optimization algorithm five times per problem. We set the maximum computational time to 24 h for each run.

The results of the computational experiments are summarized in **Table 4**, and the convergence curves are shown in **Figure 7**. UNDX/MGG failed to find solutions for all problems. MATLAB GA found solutions for B5 and B6. eSS and $REX^{star}$/JGG found solutions for 5 out of 6 problems. eSS failed in the HIV problem, and $REX^{star}$/JGG failed in B4. In summary, eSS and $REX^{star}$/JGG outperformed MATLAB GA and UNDX/MGG. Notably, only $REX^{star}$/JGG found solutions in the HIV problem, and only eSS found solutions in the B4 problem. In the directory RCGAToolbox/doc/benchmark/bio and RCGAToolbox/doc/benchmark/biopredyn-bench, the scripts for the computational experiment are provided. Neither IQM Tools nor SundialsTB is required for the experiments. Please note that the optimization algorithms have several options, and we kept them as default as possible in the computational experiments.

#### 7.5. Speed-up by parallel computation

We investigated how much RCGAs could be accelerated by parallel computation. We varied the number of computation cores (i.e., the degree of parallelization) and calculated the speed-up factor:

$$\begin{aligned} \textit{Speed-up Factor} &= \frac{\textit{Computational Speed for Parallel Computation}}{\textit{Computational Speed for Sequential Computation}} \\ &= \frac{\textit{Computational Time for Sequential Computation}}{\textit{Computational Time for Parallel Computation}} \end{aligned} \quad . \ldots\ldots\ldots\ldots (6)$$

As the number of computational cores increased, the speed-up factor increased, except in the case of 32 cores in the three-step problem (**Figure 8**). The speed-up factor was maximized in the three-step problem when 16 cores were used: 6.0 ± 0.2 for UNDX/MGG and 4.3 ± 0.1 for $\mathrm{REX}^{\mathrm{star}}$/JGG. In the HIV problem, the speed-up factor was maximized when 32 cores were used: 12.0 ± 0.3 for UNDX/MGG and 7.4 ± 0.6 for $\mathrm{REX}^{\mathrm{star}}$/JGG.

It should be noted that, in general, the ideal linear speed-up (black dashed lines in **Figure 8**) is rarely achieved because (i) some parts in programs cannot be processed in parallel, and (ii) the cost of intercore communication increases as the number of cores increases. In the three-step problem, the speed-up factor for 32 cores is slightly lower than that for 16 cores, probably because of an increased communication cost. The maximum speed-up factor for the three-step problem is lower than that for the HIV problem, because the ODEs for the three-step model can be very stiff depending on the model parameters, and thus parallelization does not help much. Since quad-core CPUs are common, the speed-up factor for four cores is practically important. In the three-step problem, the speed-up factor for four cores is 3.1 ± 0.1 for UNDX/MGG and 2.8 ± 0.02 for $\mathrm{REX}^{\mathrm{star}}$/JGG. In the HIV problem, it is 3.5 ± 0.02 for UNDX/MGG and 3.1 ± 0.1 for $\mathrm{REX}^{\mathrm{star}}$/JGG. In summary, parallel computation accelerates parameter estimation in kinetic modeling. In the directory RCGAToolbox/doc/benchmark/parallel, scripts for the computational experiment are provided.

#### Appendix A. Installation of optional toolboxes

This appendix explains how to install the optional toolboxes: IQM Tools, SundialsTB, and libSBML. If the RCGAToolbox does not work properly, run the diagnosis script **RCGAToolbox_Diagnosis.m** to verify that the optional toolboxes are installed. If SBML-related functions or the fast ODE solver option (**fast_flag** = 2) are not available, it is likely that IQM Tools or libSBML are not properly installed. If the fast ODE solver option (**fast_flag** = 1) is not available, it is likely that SundialsTB is not properly installed. For the installation of these optional toolboxes, please follow their respective installation notes. For those who are unfamiliar with these toolboxes, we provide short installation notes below. We successfully installed IQM Tools, SundialsTB, and libSBML in the following environments: (i) Windows 10 (2004) with MATLAB R2016a, (ii) SUSE Linux Enterprise Server 11 (x86_64) with MATLAB R2016a, and (iii) macOS Big Sur (11.1, Intel CPU) with MATLAB R2020b.

##### A.1. IQM Tools

1. Make sure that you can use the MEX command on MATLAB.
2. Download IQM Tools (IQMtools V1.2.2.2.zip) from the IntiQuan website: https://iqmtools.intiquan.com/.
3. Unzip it in the directory where you want to install it (typically, Documents/MATLAB).
4. Run installIQMtoolsInitial.m. Depending on the C compilers, errors may occur while compiling IQM Tools Pro. In that case, run installIQMpro.m and modify CVODEmex25.c according to the error messages. We found that only minor modifications were required.
5. After running installIQMtoolsInitial.m successfully, save the paths on MATLAB if you prefer (or run installIQMtools.m each time you start MATLAB).

##### A.2. SundialsTB

1. Make sure that you can use the MEX command on MATLAB.
2. Download Sundials (sundials-2.6.2.tar.gz) from the Lawrence Livermore National Laboratory website: https://computing.llnl.gov/projects/sundials. SundialsTB is included in the (earlier) Sundials packages.
3. Decompress the Sundials package in the directory where you want to install it (typically, Documents/MATLAB).
4. Run install_STB.m in the SundialsTB directory. Answer “y” for questions about MEX options and the CVODES interface and “n” for others. If asked, name the install directory, or leave it blank.
5. Run startup_STB.m with the path to the install directory as an input (typically, “startup_STB(‘.’)” without extension under the installation directory).
6. Save the paths in MATLAB if you prefer (or run startup_STB.m every time you start MATLAB).

##### A.3. libSBML

IQM Tools includes libSBML for Windows but not Linux or macOS. To use IQM Tools on Linux or macOS, libSBML must be installed separately.

1. Download libSBML for MATLAB (libSBML-5.18.0-matlab-binaries.tar.gz) from SourceForge: https://sourceforge.net/projects/sbml/files/libsbml/MATLAB%20Interface/.
2. Decompress it in the directory where you want to install it (typically, Documents/MATLAB/libSBML).
3. On MATLAB, save the path to the directory where you have libSBML binaries (typically, Documents/MATLAB/libSBML).
4. macOS may automatically block libSBML. In that case, you need to approve libSBML from the Security & Privacy Panel.

### Appendix B. List of RCGAToolbox functions

To use the functions provided in the RCGAToolbox, type "help *function_name*" in the MATLAB Command Window. The RCGAToolbox has the functions shown below.

#### B.1. RCGAToolbox/source/RCGA

RCGAToolbox/source/RCGA/shared

- RCGA_Main

RCGAToolbox/source/RCGA/shared/sort

- RCGAsrsort
- RCGAswap

RCGAToolbox/source/RCGA/shared/IO

- RCGAprintTransition
- RCGAprintWelcomeMessage
- RCGAwriteBest
- RCGAwritePopulation
- RCGAwriteTransition

RCGAToolbox/source/RCGA/shared/misc

- RCGAcheckInputs
- RCGAfindBest
- RCGAgetFitness
- RCGAgetInitPopulation
- RCGArequestFitnessCalc
- ScriptInitTransition
- ScriptStoreBestAndFinalPopulation
- ScriptStoreTransition

RCGAToolbox/source/RCGA/shared/local

- cst_wrapper
- obj_wrapper
- RCGAlocalOptimize

RCGAToolbox/source/RCGA/UNDXMGG

- RCGA_MGG
- RCGA_UNDX
- RCGA_UNDXMGG
- RCGAgetNewChild

RCGAToolbox/source/RCGA/REXstarJGG

- RCGA_JGG
- RCGA_REXstar
- RCGA_REXstarJGG

RCGAToolbox/source/RCGA/misc

- RCGA_CustomRCGA
- RCGAdefaultfinalreportfun
- RCGAdefaultinterimreportfun

#### B.2. RCGAToolbox/source/PE

- RCGA_PE
- RCGA_REXstarJGG_PE
- RCGA_UNDXMGG_PE

RCGAToolbox/source/PE/sim

- odestb
- RCGAsimulate
- RCGAsimulateMEX
- RCGAsimulateODE
- RCGAsimulateSTB
- wrapper_odefun

RCGAToolbox/source/PE/misc

- RCGAcreateConciseODEfile
- RCGAcreateODEfile
- RCGAmakeMEXmodel
- RCGAplotter
- RCGAreadConciseODEfile
- RCGAreplaceWords
- RCGAssr

RCGAToolbox/source/PE/IO

- RCGAinterimreportfun_PE

#### B.3. RCGAToolbox/source/GUI

- RCGAToolbox_MissionControl
- RCGAToolbox_MissionControl_PE
- RCGAcreateExecutableScript
- RCGAcreateExecutableScript_PE

#### Appendix C. Unconstrained test problems

We employed 14 unconstrained test problems: 13 from previous studies (Akimoto, et al., 2009; Kobayashi, 2009; Maeda, et al., 2018; Maeda and Kurata, 2009) and an additional problem (the scaled shifted rotated Rastrign). All the unconstrained test problems except Sphere have one or more properties known to hamper the search: ill-scale, intervariable dependency, and multimodality (Kobayashi, 2009). In ill-scaled problems, some decision variables significantly impact the objective function $f$, but others have only a small impact. Intervariable-dependent problems cannot be divided into smaller problems. Thus, in intervariable dependent problems, multiple decision variables need to be simultaneously tuned to reduce $f$. Multimodal problems have multiple local minima.

##### C.1. Sphere

Minimize:

$$f(\mathbf{x}) = \sum_{i=1}^{n} x_i^2 \quad \text{(C1a)}$$

Subject to:

$$-5.12 \le x_i \le 5.12 \quad (i = 1, ..., n) \quad \text{(C1b)}$$

The optimum is located at $\mathbf{x}^* = (0, \ldots, 0)$, where $f(\mathbf{x}^*) = 0$. $n = 100$ in our study.

##### C.2. Scaled Sphere

Minimize:

$$f(\mathbf{x}) = \sum_{i=1}^{n} (i \cdot x_i)^2 \quad \text{(C2a)}$$

Subject to:

$$-5.12 \le x_i \le 5.12 \quad (i = 1, ..., n) \quad \text{(C2b)}$$

The optimum is located at $\mathbf{x}^* = (0, \ldots, 0)$, where $f(\mathbf{x}^*) = 0$. $n = 100$ in our study. The scaled Sphere is an ill-scaled problem.

##### C.3. Ellipsoid

Minimize:

$$f(\mathbf{x}) = \sum_{i=1}^{n} (1000^{\frac{i-1}{n-1}} \cdot x_i)^2 \quad \text{(C3a)}$$

Subject to:

$$-5.12 \le x_i \le 5.12 \quad (i = 1, ..., n) \quad \text{(C3b)}$$

The optimum is located at $\mathbf{x}^* = (0, \ldots, 0)$, where $f(\mathbf{x}^*) = 0$. $n = 100$ in our study. Ellipsoid is an ill-scaled problem.

#### C.4. Cigar

Minimize:

$$f(\mathbf{x}) = x_1^2 + \sum_{i=2}^{n} (1000 \cdot x_i)^2 \quad \text{(C4a)}$$

Subject to:

$$-5.12 \le x_i \le 5.12 \quad (i = 1, ..., n) \quad \text{(C4b)}$$

The optimum is located at $\mathbf{x}^* = (0, \ldots, 0)$, where $f(\mathbf{x}^*) = 0$. $n = 100$ in our study. Cigar is an ill-scaled problem.

#### C.5. *k*-Tablet

Minimize:

$$f(\mathbf{x}) = \sum_{i=1}^{k} x_i^2 + \sum_{i=k+1}^{n} (100 \cdot x_i)^2 \quad \text{(C5a)}$$

Subject to:

$$-5.12 \le x_i \le 5.12 \quad (i = 1, ..., n), \quad \text{(C5b)}$$

where $k = n/4$. The optimum is located at $\mathbf{x}^* = (0, \ldots, 0)$, where $f(\mathbf{x}^*) = 0$. $n = 100$ in our study. The *k*-Tablet is an ill-scaled problem.

#### C.6. MM benchmark

Minimize:

$$f(\mathbf{x}) = \sum_{i=1}^{n/2} \left[ \left( \frac{10^{x_{2i-1}} \cdot 0.1}{10^{x_{2i}-2} + 0.1} - 3.33 \right)^2 + \left( \frac{10^{x_{2i-1}} \cdot 5.0}{10^{x_{2i}-2} + 5.0} - 4.95 \right)^2 \right] \quad \text{(C6a)}$$

Subject to:

$$-1 \le x_i \le 1 \quad (i = 1, ..., n) \quad \text{(C6b)}$$

The optimum is located at $\mathbf{x}^* = (0.699, \ldots, 0.699)$, where $f(\mathbf{x}^*) = 0$. $n = 100$ in our study. The MM benchmark is a bio-inspired benchmark problem: $V_{max}$ ($10^{x_{2i-1}}$) and $K_m$ ($10^{x_{2i}-2}$) in a saturated Michaelis-Menten rate equation are estimated based on two given points: the substrate level and reaction rate are 0.1 and 3.33 for one data point and 5.0 and 4.95 for another data point. The MM benchmark is an ill-scaled and intervariable-dependent problem.

#### C.7. Rosenbrock (star)

Minimize:

$$f(\mathbf{x}) = \sum_{i=2}^{n} \left[ 100(x_1 - x_i^2)^2 + (1 - x_i)^2 \right] \quad \text{(C7a)}$$

Subject to:

$$-2.048 \le x_i \le 2.048 \quad (i = 1, ..., n) \quad \text{(C7b)}$$

The optimum is located at $\mathbf{x}^* = (1, \ldots, 1)$, where $f(\mathbf{x}^*) = 0$. $n = 100$ in our study. Rosenbrock (star) is an intervariable-dependent problem.

#### C.8. Rosenbrock (chain)

Minimize:

$$f(\mathbf{x}) = \sum_{i=1}^{n-1} \left[ 100(x_{i+1} - x_i^2)^2 + (1 - x_i)^2 \right] \quad \text{(C8a)}$$

Subject to:

$$-2.048 \le x_i \le 2.048 \quad (i = 1, ..., n) \quad \text{(C8b)}$$

The optimum is located at $\mathbf{x}^* = (1, \ldots, 1)$, where $f(\mathbf{x}^*) = 0$. $n = 100$ in our study. Rosenbrock (chain) is an intervariable-dependent problem.

#### C.9. Ackley

Minimize:

$$f(\mathbf{x}) = 20 - 20\exp\left(-0.2\sqrt{\frac{1}{n}\sum_{i=1}^{n} x_i^2}\right) + e - \exp\left(\frac{1}{n}\sum_{i=1}^{n}\cos(2\pi x_i)\right) \quad \text{(C9a)}$$

Subject to:

$$-32.768 \le x_i \le 32.768 \quad (i = 1, ..., n) \quad \text{(C9b)}$$

The optimum is located at $\mathbf{x}^* = (0, \ldots, 0)$, where $f(\mathbf{x}^*) = 0$. $n = 100$ in our study. Ackley is an intervariable-dependent and multimodal problem.

##### C.10. Bohachevsky

Minimize:

$$f(\mathbf{x}) = \sum_{i=1}^{n-1} \left[ x_i^2 + 2x_{i+1}^2 - 0.3\cos(3\pi x_i) - 0.4\cos(4\pi x_{i+1}) + 0.7 \right] \quad \text{(C10a)}$$

Subject to:

$$-5.12 \le x_i \le 5.12 \quad (i = 1, ..., n) \quad \text{(C10b)}$$

The optimum is located at $\mathbf{x}^* = (0, \ldots, 0)$, where $f(\mathbf{x}^*) = 0$. $n = 100$ in our study. Bohachevsky is a multimodal problem.

##### C.11. Rastrigin

Minimize:

$$f(\mathbf{x}) = 10n + \sum_{i=1}^{n} \left[ x_i^2 - 10\cos(2\pi x_i) \right] \quad \text{(C11a)}$$

Subject to:

$$-5.12 \le x_i \le 5.12 \quad (i = 1, ..., n) \quad \text{(C11b)}$$

The optimum is located at $\mathbf{x}^* = (0, \ldots, 0)$, where $f(\mathbf{x}^*) = 0$. $n = 50$ in our study. Rastrigin is a multimodal problem.

##### C.12. Schaffer

Minimize:

$$f(\mathbf{x}) = \sum_{i=1}^{n-1} \left[ (x_i^2 + x_{i+1}^2)^{0.25} \cdot \left[ \sin^2(50(x_i^2 + x_{i+1}^2)^{0.1}) + 1.0 \right] \right] \quad \text{(C12a)}$$

Subject to:

$$-100 \le x_i \le 100 \quad (i = 1, ..., n) \quad \text{(C12b)}$$

The optimum is located at $\mathbf{x}^* = (0, \ldots, 0)$, where $f(\mathbf{x}^*) = 0$. $n = 50$ in our study. Schaffer is an intervariable-dependent and multimodal problem.

##### C.13. Schwefel

Minimize:

$$f(\mathbf{x}) = 418.9828873n + \sum_{i=1}^{n} x_i \sin\sqrt{|x_i|} \quad \text{(C13a)}$$

Subject to:

$$-512 \le x_i \le 512 \quad (i = 1, ..., n) \quad \text{(C13b)}$$

The optimum is located at $\mathbf{x}^* = (-420.9687, \ldots, -420.9687)$, where $f(\mathbf{x}^*) = 0$. $n = 20$ in our study. Schwefel is a multimodal problem.

##### C.14. Scaled shifted rotated Rastrigin

Minimize:

$$f(\mathbf{z}) = 10n + \sum_{i=1}^{n} \left[ z_i^2 - 10\cos(2\pi z_i) \right], \quad \text{(C14a)}$$

$$\mathbf{z} = \mathbf{R} \cdot \mathbf{y}, \quad \text{(C14b)}$$

$$\mathbf{y} = \mathbf{M} \cdot \mathbf{x} - 4. \quad \text{(C14c)}$$

Subject to:

$$-5.12 \le x_i \le 5.12 \quad (i = 1, ..., n) \quad \text{(C14d)}$$

$\mathbf{R}$ is an $n \times n$ rotation matrix, and $\mathbf{M}$ is the diagonal matrix represented by diag(1, 2, ..., $n$). The optimum is located at $\mathbf{x}^* = (4, 2, \ldots, 4/n)$, where $f(\mathbf{x}^*) = 0$. $n = 25$ in our study. The scaled shifted rotated Rastrigin is an ill-scaled, intervariable-dependent, and multimodal problem.

#### Appendix D. Constrained test problems

We adopted 13 constrained test problems from previous studies (Ji and Xu, 2006; Koziel and Michalewicz, 1999; Runarsson and Yao, 2000; Runarsson and Yao, 2005). We transformed the maximization problems to minimization problems and equality constraints to inequality constraints with allowable error ($\delta$).

### D.1. g01

Minimize:

$$f(\mathbf{x}) = 5\sum_{i=1}^{4} x_i - 5\sum_{i=1}^{4} x_i^2 - \sum_{i=5}^{13} x_i \quad \text{(D1a)}$$

Subject to:

$$\begin{aligned}
g_1(\mathbf{x}) &= 2x_1 + 2x_2 + x_{10} + x_{11} - 10 \le 0 \\
g_2(\mathbf{x}) &= 2x_1 + 2x_3 + x_{10} + x_{12} - 10 \le 0 \\
g_3(\mathbf{x}) &= 2x_2 + 2x_3 + x_{11} + x_{12} - 10 \le 0 \\
g_4(\mathbf{x}) &= -8x_1 + x_{10} \le 0 \\
g_5(\mathbf{x}) &= -8x_2 + x_{11} \le 0 \\
g_6(\mathbf{x}) &= -8x_3 + x_{12} \le 0 \\
g_7(\mathbf{x}) &= -2x_4 - x_5 + x_{10} \le 0 \\
g_8(\mathbf{x}) &= -2x_6 - x_7 + x_{11} \le 0 \\
g_9(\mathbf{x}) &= -2x_8 - x_9 + x_{12} \le 0
\end{aligned} \quad \text{(D1b)}$$

$$\begin{aligned}
&0 \le x_i \le 1 \quad (i = 1, \ldots, 9) \\
&0 \le x_i \le 100 \quad (i = 10, 11, 12) \\
&0 \le x_{13} \le 1
\end{aligned} \quad \text{(D1c)}$$

The optimum is located at $\mathbf{x}^* = (1, 1, 1, 1, 1, 1, 1, 1, 1, 3, 3, 3, 1)$, where $f(\mathbf{x}^*) = -15$.

**D.2. g02**

Minimize:

$$f(\mathbf{x}) = -\left| \frac{\sum_{i=1}^{n} \cos^4(x_i) - 2\prod_{i=1}^{n} \cos^2(x_i)}{\sqrt{\sum_{i=1}^{n} \left(i \cdot x_i^2\right)}} \right| \quad \text{(D2a)}$$

Subject to:

$$g_1(\mathbf{x}) = 0.75 - \prod_{i=1}^{n} x_i \le 0$$

$$g_2(\mathbf{x}) = \sum_{i=1}^{n} x_i - 7.5n \le 0 \quad \text{(D2b)}$$

$$0 \le x_i \le 10 \quad (i = 1, \ldots, n), \quad \text{(D2c)}$$

where $n$ = 20. The optimum is unknown. The best known value is $f(\mathbf{x}^*)$ = -0.803619.

**D.3. g03**

Minimize:

$$f(\mathbf{x}) = -\left(\sqrt{n}\right)^n \prod_{i=1}^{n} x_i \quad \text{(D3a)}$$

Subject to:

$$g_1(\mathbf{x}) = \left| \sum_{i=1}^{n} x_i^2 - 1 \right| - \delta \le 0 \quad \text{(D3b)}$$

$$0 \le x_i \le 1 \quad (i = 1, \ldots, n), \quad \text{(D3c)}$$

where $n$ = 10 and $\delta = 10^{-4}$. The optimum is located at $\mathbf{x}^* = (1/\sqrt{n},\ 1/\sqrt{n},\ ...)$, where $f(\mathbf{x}^*)$ = -1.

## D.4. g04

Minimize:

$$f(\mathbf{x}) = 5.3578547x_3^2 + 0.8356891x_1x_5 + 37.293239x_1 - 40792.141 \quad \text{(D4a)}$$

Subject to:

$$\begin{aligned}
g_1(\mathbf{x}) &= 85.334407 + 0.0056858x_2x_5 + 0.0006262x_1x_4 - 0.0022053x_3x_5 - 92 \le 0 \\
g_2(\mathbf{x}) &= -85.334407 - 0.0056858x_2x_5 - 0.0006262x_1x_4 + 0.0022053x_3x_5 \le 0 \\
g_3(\mathbf{x}) &= 80.51249 + 0.0071317x_2x_5 + 0.0029955x_1x_2 + 0.0021813x_3^2 - 110 \le 0 \\
g_4(\mathbf{x}) &= -80.51249 - 0.0071317x_2x_5 - 0.0029955x_1x_2 - 0.0021813x_3^2 + 90 \le 0 \\
g_5(\mathbf{x}) &= 9.300961 + 0.0047026x_3x_5 + 0.0012547x_1x_3 + 0.0019085x_3x_4 - 25 \le 0 \\
g_6(\mathbf{x}) &= -9.300961 - 0.0047026x_3x_5 - 0.0012547x_1x_3 - 0.0019085x_3x_4 + 20 \le 0
\end{aligned} \quad \text{(D4b)}$$

$$\begin{aligned}
&78 \le x_1 \le 102 \\
&33 \le x_2 \le 45 \\
&27 \le x_i \le 45 \quad (i = 3,\ 4,\ 5)
\end{aligned} \quad \text{(D4c)}$$

The optimum is located at $\mathbf{x}^*$= (78, 33, 29.995256025682, 45, 36.775812905788), where $f(\mathbf{x}^*)$ = -30665.539.

## D.5. g05

Minimize:

$$f(\mathbf{x}) = 3x_1 + 0.000001x_1^3 + 2x_2 + (0.000002/3)x_2^3 \quad \text{(D5a)}$$

Subject to:

$$\begin{aligned}
g_1(\mathbf{x}) &= -x_4 + x_3 - 0.55 \le 0 \\
g_2(\mathbf{x}) &= -x_3 + x_4 - 0.55 \le 0 \\
g_3(\mathbf{x}) &= |1000\sin(-x_3 - 0.25) + 1000\sin(-x_4 - 0.25) + 894.8 - x_1| - \delta \le 0 \\
g_4(\mathbf{x}) &= |1000\sin(x_3 - 0.25) + 1000\sin(x_3 - x_4 - 0.25) + 894.8 - x_2| - \delta \le 0 \\
g_5(\mathbf{x}) &= |1000\sin(x_4 - 0.25) + 1000\sin(x_4 - x_3 - 0.25) + 1294.8| - \delta \le 0
\end{aligned} \quad \text{(D5b)}$$

$$\begin{aligned}
&0 \le x_i \le 1200 \quad (i = 1, 2) \\
&-0.55 \le x_i \le 0.55 \quad (i = 3, 4)
\end{aligned}, \quad \text{(D5c)}$$

where $\delta = 10^{-4}$. The known best value is $\mathbf{x}^*$ = (679.9453, 1026.067, 0.1188764, -0.3962336), where $f(\mathbf{x}^*)$ = 5126.4981.

## D.6. g06

Minimize:

$$f(\mathbf{x}) = (x_1 - 10)^3 + (x_2 - 20)^3 \quad \text{(D6a)}$$

Subject to:

$$\begin{aligned} g_1(\mathbf{x}) &= -(x_1 - 5)^2 - (x_2 - 5)^2 + 100 \le 0 \\ g_2(\mathbf{x}) &= (x_1 - 6)^2 + (x_2 - 5)^2 - 82.81 \le 0 \end{aligned} \quad \text{(D6b)}$$

$$\begin{aligned} 13 \le x_1 \le 100 \\ 0 \le x_2 \le 100 \end{aligned} \quad \text{(D6c)}$$

The optimum is located at $\mathbf{x}^*$ = (14.095, 0.84296), where $f(\mathbf{x}^*)$ = -6961.81388.

## D.7. g07

Minimize:

$$\begin{aligned} f(\mathbf{x}) = {} & x_1^2 + x_2^2 + x_1x_2 - 14x_1 - 16x_2 + (x_3 - 10)^2 + 4(x_4 - 5)^2 + (x_5 - 3)^2 \\ & + 2(x_6 - 1)^2 + 5x_7^2 + 7(x_8 - 11)^2 + 2(x_9 - 10)^2 + (x_{10} - 7)^2 + 45 \end{aligned} \quad \text{(D7a)}$$

Subject to:

$$\begin{aligned} g_1(\mathbf{x}) &= -105 + 4x_1 + 5x_2 - 3x_7 + 9x_8 \le 0 \\ g_2(\mathbf{x}) &= 10x_1 - 8x_2 - 17x_7 + 2x_8 \le 0 \\ g_3(\mathbf{x}) &= -8x_1 + 2x_2 + 5x_9 - 2x_{10} - 12 \le 0 \\ g_4(\mathbf{x}) &= 3(x_1 - 2)^2 + 4(x_2 - 3)^2 + 2x_3^2 - 7x_4 - 120 \le 0 \\ g_5(\mathbf{x}) &= 5x_1^2 + 8x_2 + (x_3 - 6)^2 - 2x_4 - 40 \le 0 \\ g_6(\mathbf{x}) &= x_1^2 + 2(x_2 - 2)^2 - 2x_1x_2 + 14x_5 - 6x_6 \le 0 \\ g_7(\mathbf{x}) &= 0.5(x_1 - 8)^2 + 2(x_2 - 4)^2 + 3x_5^2 - x_6 - 30 \le 0 \\ g_8(\mathbf{x}) &= -3x_1 + 6x_2 + 12(x_9 - 8)^2 - 7x_{10} \le 0 \end{aligned} \quad \text{(D7b)}$$

$$-10 \le x_i \le 10 \quad (i = 1, \ldots, 10) \quad \text{(D7c)}$$

The optimum is located at $\mathbf{x}^*$ = (2.171996, 2.363683, 8.773926, 5.095984, 0.9906548, 1.430574, 1.321644, 9.828726, 8.280092, 8.375927), where $f(\mathbf{x}^*)$ = 24.3062091.

**D.8. g08**

Minimize:

$$f(\mathbf{x}) = -\frac{\sin^3(2\pi x_1)\cdot\sin(2\pi x_2)}{x_1^3(x_1 + x_2)} \quad \text{(D8a)}$$

Subject to:

$$\begin{aligned} g_1(\mathbf{x}) &= x_1^2 - x_2 + 1 \le 0 \\ g_2(\mathbf{x}) &= 1 - x_1 + (x_2 - 4)^2 \le 0 \end{aligned} \quad \text{(D8b)}$$

$$0 \le x_i \le 10 \quad (i = 1, 2) \quad \text{(D8c)}$$

The optimum is located at $\mathbf{x}^*$ = (1.2279713, 4.2453733), where $f(\mathbf{x}^*)$ = -0.095825.

**D.9. g09**

Minimize:

$$\begin{aligned} f(\mathbf{x}) = &(x_1 - 10)^2 + 5(x_2 - 12)^2 + x_3^4 + 3(x_4 - 11)^2 \\ &+ 10x_5^6 + 7x_6^2 + x_7^4 - 4x_6x_7 - 10x_6 - 8x_7 \end{aligned} \quad \text{(D9a)}$$

Subject to:

$$\begin{aligned} g_1(\mathbf{x}) &= -127 + 2x_1^2 + 3x_2^4 + x_3 + 4x_4^2 + 5x_5 \le 0 \\ g_2(\mathbf{x}) &= -282 + 7x_1 + 3x_2 + 10x_3^2 + x_4 - x_5 \le 0 \\ g_3(\mathbf{x}) &= -196 + 23x_1 + x_2^2 + 6x_6^2 - 8x_7 \le 0 \\ g_4(\mathbf{x}) &= 4x_1^2 + x_2^2 - 3x_1x_2 + 2x_3^2 + 5x_6 - 11x_7 \le 0 \end{aligned} \quad \text{(D9b)}$$

$$-10 \le x_i \le 10 \quad (i = 1, \ldots, 7) \quad \text{(D9c)}$$

The optimum is located at $\mathbf{x}^*$ = (2.330499, 1.951372, -0.4775414, 4.365726, -0.6244870, 1.038131, 1.594227), where $f(\mathbf{x}^*)$ = 680.6300573.

### D.10. g10

Minimize:

$$f(\mathbf{x}) = x_1 + x_2 + x_3 \qquad \text{(D10a)}$$

Subject to:

$$\begin{aligned}
g_1(\mathbf{x}) &= -1 + 0.0025(x_4 + x_6) \le 0 \\
g_2(\mathbf{x}) &= -1 + 0.0025(x_5 + x_7 - x_4) \le 0 \\
g_3(\mathbf{x}) &= -1 + 0.01(x_8 - x_5) \le 0 \\
g_4(\mathbf{x}) &= -x_1 x_6 + 833.33252 x_4 + 100 x_1 - 83333.333 \le 0 \\
g_5(\mathbf{x}) &= -x_2 x_7 + 1250 x_5 + x_2 x_4 - 1250 x_4 \le 0 \\
g_6(\mathbf{x}) &= -x_3 x_8 + 1250000 + x_3 x_5 - 2500 x_5 \le 0
\end{aligned} \qquad \text{(D10b)}$$

$$\begin{aligned}
&100 \le x_1 \le 10000 \\
&1000 \le x_i \le 10000 \quad (i = 2,3) \\
&10 \le x_i \le 1000 \quad (i = 4,\ldots,8)
\end{aligned} \qquad \text{(D10c)}$$

The optimum is located at $\mathbf{x}^*$ = (579.3167, 1359.943, 5110.071, 182.0174, 295.5985, 217.9799, 286.4162, 395.5979), where $f(\mathbf{x}^*)$ = 7049.3307.

### D.11. g11

Minimize:

$$f(\mathbf{x}) = x_1^2 + (x_2 - 1)^2 \qquad \text{(D11a)}$$

Subject to:

$$g_1(\mathbf{x}) = \left| x_2 - x_1^2 \right| - \delta \le 0 \qquad \text{(D11b)}$$

$$-1 \le x_i \le 1 \quad (i = 1,\ 2), \qquad \text{(D11c)}$$

where $\delta = 10^{-4}$. The optimum is located at $\mathbf{x}^* = (\pm 1/\sqrt{2}, 1/2)$, where $f(\mathbf{x}^*) = 0.75$.

### D.12. g12

Minimize:

$$f(\mathbf{x}) = -\left[100-(x_1-5)^2-(x_2-5)^2-(x_3-5)^2\right]/100 \quad \text{(D12a)}$$

Subject to:

$$g_1(\mathbf{x}) = (x_1-p)^2+(x_2-q)^2+(x_3-r)^2-0.0625 \le 0, \quad \text{(D12b)}$$

$$0 \le x_i \le 10 \quad (i=1,2,3), \quad \text{(D12c)}$$

where $p$, $q$, $r$ = 1, 2, …, 9. The feasible region of the search space consists of $9^3$ disjointed spheres. A point ($x_1$, $x_2$, $x_3$) is feasible if and only if there exists $p$, $q$, $r$ such that the above inequality holds. The optimum is located at $\mathbf{x}^*$ = (5, 5, 5), where $f(\mathbf{x}^*)$ = -1.

### D.13. g13

Minimize:

$$f(\mathbf{x}) = \exp\left(\prod_{i=1}^{5} x_i\right) \quad \text{(D13a)}$$

Subject to:

$$\begin{aligned} g_1(\mathbf{x}) &= \left|\sum_{i=1}^{5} x_i^2 - 10\right| - \delta \le 0 \\ g_2(\mathbf{x}) &= \left|x_2x_3 - 5x_4x_5\right| - \delta \le 0 \\ g_3(\mathbf{x}) &= \left|x_1^3 + x_2^3 + 1\right| - \delta \le 0 \end{aligned} \quad \text{(D13b)}$$

$$\begin{aligned} -2.3 &\le x_i \le 2.3 \quad (i=1,2) \\ -3.2 &\le x_i \le 3.2 \quad (i=3,4,5) \end{aligned}, \quad \text{(D13c)}$$

where $\delta = 10^{-4}$. The optimum is located at $\mathbf{x}^*$ = (-1.717143, 1.595709, 1.827247, -0.7636413, -0.763645), where $f(\mathbf{x}^*)$ = 0.0539498.

### Appendix E. Biological test problems

#### E.1. Three-step model

The first biological test problem is the parameter estimation of the three-step model (Ji and Xu, 2006; Mendes, 2001; Moles, et al., 2003). The network map is shown in **Figure 9**. In the three-step model, substrate $S$ is converted into product $P$ through intermediates $M_1$ and $M_2$. The metabolic reactions are catalyzed by three enzymes, $E_1$, $E_2$, and $E_3$. The metabolites regulate the expression of mRNAs $G_1$, $G_2$, and $G_3$, from which the proteins $E_1$, $E_2$, and $E_3$ are translated, respectively. The values of $S$ and $P$ are fixed. Thus, the three-step model has eight variables and is described by the following ODEs:

$$\frac{dG_1}{dt} = \frac{V_1}{1 + (P/Ki_1)^{ni_1} + (Ka_1/S)^{na_1}} - k_1 \cdot G_1$$

$$\frac{dG_2}{dt} = \frac{V_2}{1 + (P/Ki_2)^{ni_2} + (Ka_2/M_1)^{na_2}} - k_2 \cdot G_2$$

$$\frac{dG_3}{dt} = \frac{V_3}{1 + (P/Ki_3)^{ni_3} + (Ka_3/M_2)^{na_3}} - k_3 \cdot G_3$$

$$\frac{dE_1}{dt} = \frac{V_4 \cdot G_1}{K_4 + G_1} - k_4 \cdot E_1$$

$$\frac{dE_2}{dt} = \frac{V_5 \cdot G_2}{K_5 + G_2} - k_5 \cdot E_2$$

$$\frac{dE_3}{dt} = \frac{V_6 \cdot G_3}{K_6 + G_3} - k_6 \cdot E_3$$

$$\frac{dM_1}{dt} = \frac{kcat_1 \cdot E_1 \cdot (S - M_1)/Km_1}{1 + S/Km_1 + M_1/Km_2} - \frac{kcat_2 \cdot E_2 \cdot (M_1 - M_2)/Km_3}{1 + M_1/Km_3 + M_2/Km_4}$$

$$\frac{dM_2}{dt} = \frac{kcat_2 \cdot E_2 \cdot (M_1 - M_2)/Km_3}{1 + M_1/Km_3 + M_2/Km_4} - \frac{kcat_3 \cdot E_3 \cdot (M_2 - P)/Km_5}{1 + M_2/Km_5 + P/Km_6} \quad \text{..........................(E1)}$$

The initial values are $G_1 = 0.66667$, $G_2 = 0.57254$, $G_3 = 0.41758$, $E_1 = 0.4$, $E_2 = 0.36409$, $E_3 = 0.29457$, $M_1 = 1.419$, and $M_2 = 0.93464$. The three-step model has 36 kinetic parameters to be estimated (**Table 5**). The parameter estimation problem is the minimization of the sum of squared differences between the pseudo-experimental values and the simulated values:

Minimize:

$$f(\mathbf{x}) = \sum_{i=1}^{n_{exp}} \sum_{j=1}^{n_{point}} \sum_{k=1}^{n_{var}} (y_{i,j,k}^{sim} - y_{i,j,k}^{exp})^2 \quad \text{(E2a)}$$

Subject to:

$$\begin{array}{ll}
g_1(\mathbf{x}) = |\log_2(V_2/V_1)| - 1 \le 0 & g_{13}(\mathbf{x}) = |\log_2(V_5/V_4)| - 1 \le 0 \\
g_2(\mathbf{x}) = |\log_2(V_3/V_1)| - 1 \le 0 & g_{14}(\mathbf{x}) = |\log_2(V_6/V_4)| - 1 \le 0 \\
g_3(\mathbf{x}) = |\log_2(Ki_2/Ki_1)| - 1 \le 0 & g_{15}(\mathbf{x}) = |\log_2(K_5/K_4)| - 1 \le 0 \\
g_4(\mathbf{x}) = |\log_2(Ki_3/Ki_1)| - 1 \le 0 & g_{16}(\mathbf{x}) = |\log_2(K_6/K_4)| - 1 \le 0 \\
g_5(\mathbf{x}) = |\log_2(ni_2/ni_1)| - 1 \le 0 & g_{17}(\mathbf{x}) = |\log_2(k_5/k_4)| - 1 \le 0 \\
g_6(\mathbf{x}) = |\log_2(ni_3/ni_1)| - 1 \le 0 & g_{18}(\mathbf{x}) = |\log_2(k_6/k_4)| - 1 \le 0 \\
g_7(\mathbf{x}) = |\log_2(Ka_2/Ka_1)| - 1 \le 0 & g_{19}(\mathbf{x}) = |\log_2(kcat_2/kcat_1)| - 1 \le 0 \\
g_8(\mathbf{x}) = |\log_2(Ka_3/Ka_1)| - 1 \le 0 & g_{20}(\mathbf{x}) = |\log_2(kcat_3/kcat_1)| - 1 \le 0 \\
g_9(\mathbf{x}) = |\log_2(na_2/na_1)| - 1 \le 0 & g_{21}(\mathbf{x}) = |\log_2(Km_3/Km_1)| - 1 \le 0 \\
g_{10}(\mathbf{x}) = |\log_2(na_3/na_1)| - 1 \le 0 & g_{22}(\mathbf{x}) = |\log_2(Km_5/Km_1)| - 1 \le 0 \\
g_{11}(\mathbf{x}) = |\log_2(k_2/k_1)| - 1 \le 0 & g_{23}(\mathbf{x}) = |\log_2(Km_4/Km_2)| - 1 \le 0 \\
g_{12}(\mathbf{x}) = |\log_2(k_3/k_1)| - 1 \le 0 & g_{24}(\mathbf{x}) = |\log_2(Km_6/Km_2)| - 1 \le 0
\end{array}, \quad \text{(E2b)}$$

where $y_{i,j,k}^{exp}$ indicates the pseudo-experimental data of the $k^{th}$ variable at the $j^{th}$ data point in the $i^{th}$ experiment. $y_{i,j,k}^{sim}$ indicates the simulated data. $n_{var}$, $n_{point}$, and $n_{exp}$ are the number of model variables, number of data points, and number of pseudo-experiments, respectively. The constraint functions $g_1$-$g_{24}$ are employed to incorporate prior information such that $V_1$ – $V_3$ have similar values, $Ki_1$ – $Ki_3$ have similar values, and so on. A total of 16 pseudo-experimental data were generated by numerically integrating Eqs. (E1) using the optimum parameter values (**Table 5**) and different $S$ and $P$ concentrations (**Table 6**).

#### E.2. HIV model

The second biological test problem is the parameter estimation of the HIV model, which describes the mechanism of irreversible inhibition of HIV proteinase (Ji and Xu, 2006; Kuzmic, 1996; Mendes and Kell, 1998). The network map is shown in **Figure 10**. The enzyme is inactive in the monomeric form $M$ and active in the dimer form $E$. Product $P$ is a competitive inhibitor of substrate $S$. $I$ is an irreversible inhibitor. The HIV model has nine variables and is described by the following ODEs:

$$\frac{dM}{dt} = -2 \cdot kmd \cdot M^2 + 2 \cdot kdm \cdot E$$

$$\frac{dP}{dt} = kcat \cdot ES - kon \cdot P \cdot E + kp \cdot EP$$

$$\frac{dS}{dt} = -kon \cdot S \cdot E + ks \cdot ES$$

$$\frac{dI}{dt} = -kon \cdot I \cdot E + ki \cdot EI$$

$$\frac{dES}{dt} = kon \cdot S \cdot E - ks \cdot ES - kcat \cdot ES$$

$$\frac{dEP}{dt} = kon \cdot P \cdot E - kp \cdot EP$$

$$\frac{dE}{dt} = kmd \cdot M^2 - kdm \cdot E - kon \cdot S \cdot E + ks \cdot ES + kcat \cdot ES - kon \cdot P \cdot E + kp \cdot EP - kon \cdot I \cdot E + ki \cdot EI$$

$$\frac{dEI}{dt} = kon \cdot I \cdot E - ki \cdot EI - kde \cdot EI$$

$$\frac{dEJ}{dt} = kde \cdot EI \qquad \text{.......................................(E3)}$$

The initial values for *S*, *E*, and *I*, denoted as $S_0$, $E_0$, and $I_0$, respectively, depend on the experiments. $S_0$ and $E_0$ are to be estimated in the parameter estimation problem. $I_0$ is 0 μM for experiment A, 0.0015 μM for experiment B, 0.003 μM for experiment C, and 0.004 μM for experiments D and E. The initial values for *M*, *P*, *ES*, *EP*, *EI*, and *EJ* were zero. The following four parameters are known: *kmd* = 0.1 $\mu M^{-1}$ $s^{-1}$, *kdm* = 0.001 $s^{-1}$, *kon* = 100 $\mu M^{-1}$ $s^{-1}$, and $\varepsilon = 0.024$. In total, the HIV model has 20 parameters to be estimated (**Table** 7).

We obtained experimental data from a previous study (Ji and Xu, 2006). According to previous studies (Ji and Xu, 2006; Kuzmic, 1996; Mendes and Kell, 1998), the experimental data were produced from five time courses (Experiments A–E) at four different inhibitor concentrations measured fluorimetrically. The HIV proteinase ($E_0$, assay concentration 0.004 μM) was added to a solution of an irreversible inhibitor and a fluorogenic substrate ($S_0$, 25 μM). Five assays were conducted at four different concentrations of the inhibitor (0, 0.0015, 0.003, and 0.004 μM in replicate). The fluorescence changes were monitored for 1 h in each experiment.

In the HIV model, five rate constants (*ks*, *kcat*, *kp*, *ki*, and *kde*) and three other experiment-dependent parameters ($S_0$, $E_0$, and *offset*) are unknown. According to these studies (Ji and Xu, 2006; Kuzmic, 1996; Mendes and Kell, 1998), it is assumed that there is a certain degree of uncertainty in $S_0$ and $E_0$. Moreover, the baseline of the fluorimeter (denoted as *offset*) is not exactly zero. Thus, $S_0$, $E_0$, and *offset* need to be estimated for each experiment. The parameter estimation problem is the minimization of the sum of squared differences between the experimental and simulated values:

Minimize:

$$f(\mathbf{x}) = \sum_{i=1}^{n_{exp}} \sum_{j=1}^{n_{point}} (F_{i,j}^{sim} - F_{i,j}^{exp})^2 \quad \text{(E4a)}$$

Subject to:

$$\begin{aligned}
g_1(\mathbf{x}) &= ks - kp \le 0 \\
g_2(\mathbf{x}) &= ki - ks \le 0 \\
g_3(\mathbf{x}) &= kcat - ks \le 0 \\
g_4(\mathbf{x}) &= kde - kcat \le 0 \\
g_5(\mathbf{x}) &= \left|\log_2(S_{0,B} / S_{0,A})\right| - 1 \le 0 \\
g_6(\mathbf{x}) &= \left|\log_2(S_{0,C} / S_{0,A})\right| - 1 \le 0 \\
g_7(\mathbf{x}) &= \left|\log_2(S_{0,D} / S_{0,A})\right| - 1 \le 0 \\
g_8(\mathbf{x}) &= \left|\log_2(S_{0,E} / S_{0,A})\right| - 1 \le 0 \\
g_9(\mathbf{x}) &= \left|\log_2(E_{0,B} / E_{0,A})\right| - 1 \le 0 \\
g_{10}(\mathbf{x}) &= \left|\log_2(E_{0,C} / E_{0,A})\right| - 1 \le 0 \\
g_{11}(\mathbf{x}) &= \left|\log_2(E_{0,D} / E_{0,A})\right| - 1 \le 0 \\
g_{12}(\mathbf{x}) &= \left|\log_2(E_{0,E} / E_{0,A})\right| - 1 \le 0
\end{aligned} \quad , \quad \text{(E4b)}$$

where $F_{i,j}^{exp}$ indicates the measured fluorescence intensity of the $j^{th}$ data point in the $i^{th}$ experiment. $F_{i,j}^{sim}$ indicates the simulated fluorescence intensity. $n_{point}$ and $n_{exp}$ are the numbers of data points and experiments, respectively. $F_{i,j}^{sim}$ is calculated by

$$F_{i,j}^{sim} = \varepsilon \cdot P_{i,j}^{sim} + offset_i \, , \quad \text{(E5)}$$

where $P_{i,j}^{sim}$ indicates the simulated concentration of $P$, and $\varepsilon$ is the factor that converts the protein concentration into fluorescence intensity. The constraint functions $g_1$-$g_{12}$ are employed assuming there is some prior information about the rate constants, $S_0$, and $E_0$.

#### Tables

##### Table 1. Number of successful trials for unconstrained test problems

| Test Problem | IS | IVD | M | MATLAB GA | MEIGO eSS | RCGAToolbox UNDX/MGG | RCGAToolbox $REX^{star}$/JGG |
|---|---|---|---|---|---|---|---|
| Sphere | No | No | No | 10 | 10 | 10 | 10 |
| Scaled Sphere | Yes | No | No | 0 | 10 | 0 | 10 |
| Ellipsoid | Yes | No | No | 0 | 10 | 0 | 10 |
| Cigar | Yes | No | No | 0 | 10 | 0 | 10 |
| *k*-Tablet | Yes | No | No | 0 | 10 | 0 | 10 |
| MM Benchmark | Yes | Yes | No | 1 | 10 | 0 | 10 |
| Rosenbrock (star) | No | Yes | No | 0 | 3 | 0 | 0 |
| Rosenbrock (chain) | No | Yes | No | 0 | 6 | 0 | 10 |
| Ackley | No | Yes | Yes | 0 | 10 | 4 | 10 |
| Bohachevsky | No | No | Yes | 0 | 10 | 9 | 8 |
| Rastrigin | No | No | Yes | 6 | 10 | 6 | 1 |
| Schaffer | No | Yes | Yes | 0 | 0 | 0 | 10 |
| Schwefel | No | No | Yes | 0 | 10 | 0 | 0 |
| Scaled shifted rotated Rastrigin | Yes | Yes | Yes | 0 | 0 | 0 | 8 |

For each algorithm, 10 independent runs were performed. We employed MATLAB GA (Global Optimization Toolbox, MathWorks, Inc.) and eSS [R2019 from MEIGO (Egea, et al., 2014)] as controls. We used the RCGAToolbox for UNDX/MGG and $REX^{star}$/JGG with the local optimizer. All the unconstrained test problems except for Sphere have one or more properties that are known to hamper the search: ill-scale (IS), intervariable dependency (IVD), and multimodality (M). The computational experiment was performed on a single core of Intel Xeon E5-4650 v3 with SUSE Linux Enterprise Server 11 (x86_64).

**Table 2. Number of successful trials for constrained test problems**

| Test Problem | MATLAB GA | MEIGO eSS | RCGAToolbox UNDX/MGG | RCGAToolbox $REX^{star}$/JGG |
|---|---|---|---|---|
| g01 | 3 | 10 | 4 | 10 |
| g02 | 0 | 10 | 1 | 3 |
| g03 | 1 | 10 | 10 | 10 |
| g04 | 0 | 10 | 10 | 10 |
| g05 | 3 | 10 | 8 | 6 |
| g06 | 0 | 10 | 10 | 10 |
| g07 | 0 | 10 | 10 | 10 |
| g08 | 0 | 10 | 10 | 10 |
| g09 | 0 | 10 | 10 | 10 |
| g10 | 0 | 10 | 10 | 10 |
| g11 | 10 | 10 | 10 | 10 |
| g12 | 4 | 10 | 10 | 10 |
| g13 | 0 | 7 | 10 | 10 |

For each algorithm, 10 independent runs were performed. We employed MATLAB GA (Global Optimization Toolbox, MathWorks, Inc.) and eSS [R2019 from MEIGO (Egea, et al., 2014)] as controls. We used the RCGAToolbox for UNDX/MGG and $REX^{star}$/JGG with the local optimizer and stochastic ranking ($P_f = 0.45$). The computational experiment was performed on a single core of Intel Xeon E5-4650 v3 with SUSE Linux Enterprise Server 11 (x86_64).

**Table 3. Main features of biological test problems**

| Problem ID | Three-step | HIV | B2 | B4 | B5 | B6 |
|---|---|---|---|---|---|---|
| Reference | (Maeda, et al., 2018) | (Maeda, et al., 2018) | (Chassagnole, et al., 2002) | (Villaverde, et al., 2014) | (MacNamara, et al., 2012) | (Crombach, et al., 2012) |
| Organism | Generic | Human | *E. coli* | Chinese hamster | Generic | *D. melanogaster* |
| Description Level | Metabolism, Transcription, Translation | Protein Interaction | Metabolism | Metabolism | Signal Transduction | Developmental GRN (spatial) |
| Unknown Parameters | 36 | 20 | 116 | 117 | 86 | 37 |
| Dynamic States | 8 | 9 | 18 | 34 | 26 | 108 – 212 |
| Observed States | 8 | 1 | 9 | 13 | 6 | 108 – 212 |
| Experiments | 16 | 5 | 1 | 1 | 10 | 1 |
| Data Points | 7808 | 1500 | 110 | 169 | 960 | 1804 |
| Data Type | Simulated | Measured | Measured | Simulated | Simulated | Measured |
| Noise Level | No Noise | Real | Real | Variable | Artificial ($\sigma = 5\%$) | Real |
| Constraint Functions | 24 | 12 | 0 | 0 | 0 | 0 |
| VTR | $1.00 \times 10^{-3}$ | 0.04 | 468 | 91.4 | $6.15 \times 10^{3}$ | $2.17 \times 10^{5}$ |

**Table 4. Summary of computational experiments on biological test problems**

| Problem ID | | Three-step | HIV | B2 | B4 | B5 | B6 |
|---|---|---|---|---|---|---|---|
| VTR | | $1.00 \times 10^{-3}$ | 0.04 | 468 | 91.4 | $6.15 \times 10^{3}$ | $2.17 \times 10^{5}$ |
| MATLAB GA | Max | - | - | 866 | 17440.0 | **$6.09 \times 10^{3}$** | $4.48 \times 10^{5}$ |
| | Median | - | - | 630 | 11120.0 | **$5.37 \times 10^{3}$** | $3.84 \times 10^{5}$ |
| | Min | - | - | 538 | 3855.0 | **$3.45 \times 10^{3}$** | **$1.76 \times 10^{5}$** |
| eSS | Max | $9.97 \times 10^{-1}$ | 1.05 | **246** | 202.1 | **$3.10 \times 10^{3}$** | $2.71 \times 10^{5}$ |
| | Median | **$1.33 \times 10^{-7}$** | 0.57 | **232** | **41.0** | **$2.95 \times 10^{3}$** | **$1.56 \times 10^{5}$** |
| | Min | **$1.54 \times 10^{-8}$** | 0.56 | **230** | **35.0** | **$2.90 \times 10^{3}$** | **$1.01 \times 10^{5}$** |
| UNDX/MGG | Max | $3.97 \times 10^{2}$ | 2.28 | 1611 | 17344.2 | $1.06 \times 10^{4}$ | $2.89 \times 10^{5}$ |
| | Median | $3.97 \times 10^{2}$ | 2.07 | 1449 | 15783.4 | $9.95 \times 10^{3}$ | $2.82 \times 10^{5}$ |
| | Min | $3.97 \times 10^{2}$ | 1.48 | 1289 | 11930.9 | $8.58 \times 10^{3}$ | $2.65 \times 10^{5}$ |
| $REX^{star}$/JGG | Max | $2.14 \times 10^{0}$ | 0.56 | **253** | 279.3 | **$4.68 \times 10^{3}$** | **$9.71 \times 10^{4}$** |
| | Median | **$1.53 \times 10^{-8}$** | **0.02** | **244** | 178.3 | **$4.33 \times 10^{3}$** | **$9.62 \times 10^{4}$** |
| | Min | **$1.53 \times 10^{-8}$** | **0.02** | **222** | 168.6 | **$4.18 \times 10^{3}$** | **$9.09 \times 10^{4}$** |

The values indicate maximal (worst), median, and minimal (best) objective function values obtained in 5 trials. Bold figures indicate solutions, i.e., ≤ VTR (the value to be reached).

**Table 5. Kinetic parameters to be estimated in the three-step model**

| Parameter | Optimum | Search Space | Parameter | Optimum | Search Space |
|---|---|---|---|---|---|
| $V_1$ | 1 | $10^{-12} – 10^{6}$ | $V_4$ | 0.1 | $10^{-12} – 10^{6}$ |
| $Ki_1$ | 1 | $10^{-12} – 10^{6}$ | $K_4$ | 1 | $10^{-12} – 10^{6}$ |
| $ni_1$ | 2 | $10^{-1} – 10^{1}$ | $k_4$ | 0.1 | $10^{-12} – 10^{6}$ |
| $Ka_1$ | 1 | $10^{-12} – 10^{6}$ | $V_5$ | 0.1 | $10^{-12} – 10^{6}$ |
| $na_1$ | 2 | $10^{-1} – 10^{1}$ | $K_5$ | 1 | $10^{-12} – 10^{6}$ |
| $k_1$ | 1 | $10^{-12} – 10^{6}$ | $k_5$ | 0.1 | $10^{-12} – 10^{6}$ |
| $V_2$ | 1 | $10^{-12} – 10^{6}$ | $V_6$ | 0.1 | $10^{-12} – 10^{6}$ |
| $Ki_2$ | 1 | $10^{-12} – 10^{6}$ | $K_6$ | 1 | $10^{-12} – 10^{6}$ |
| $ni_2$ | 2 | $10^{-1} – 10^{1}$ | $k_6$ | 0.1 | $10^{-12} – 10^{6}$ |
| $Ka_2$ | 1 | $10^{-12} – 10^{6}$ | $kcat_1$ | 1 | $10^{-12} – 10^{6}$ |
| $na_2$ | 2 | $10^{-1} – 10^{1}$ | $km_1$ | 1 | $10^{-12} – 10^{6}$ |
| $k_2$ | 1 | $10^{-12} – 10^{6}$ | $km_2$ | 1 | $10^{-12} – 10^{6}$ |
| $V_3$ | 1 | $10^{-12} – 10^{6}$ | $kcat_2$ | 1 | $10^{-12} – 10^{6}$ |
| $Ki_3$ | 1 | $10^{-12} – 10^{6}$ | $km_3$ | 1 | $10^{-12} – 10^{6}$ |
| $ni_3$ | 2 | $10^{-1} – 10^{1}$ | $km_4$ | 1 | $10^{-12} – 10^{6}$ |
| $Ka_3$ | 1 | $10^{-12} – 10^{6}$ | $kcat_3$ | 1 | $10^{-12} – 10^{6}$ |
| $na_3$ | 2 | $10^{-1} – 10^{1}$ | $km_5$ | 1 | $10^{-12} – 10^{6}$ |
| $k_3$ | 1 | $10^{-12} – 10^{6}$ | $km_6$ | 1 | $10^{-12} – 10^{6}$ |

**Table 6. Substrate and product concentrations for 16 pseudo-experiments**

| Exp. | $S$ | $P$ | Exp. | $S$ | $P$ |
|---|---|---|---|---|---|
| 1 | 0.10000 | 0.05000 | 9 | 0.10000 | 0.36840 |
| 2 | 0.46416 | 0.05000 | 10 | 0.46416 | 0.36840 |
| 3 | 2.15440 | 0.05000 | 11 | 2.15440 | 0.36840 |
| 4 | 10.0000 | 0.05000 | 12 | 10.0000 | 0.36840 |
| 5 | 0.10000 | 0.13572 | 13 | 0.10000 | 1.00000 |
| 6 | 0.46416 | 0.13572 | 14 | 0.46416 | 1.00000 |
| 7 | 2.15440 | 0.13572 | 15 | 2.15440 | 1.00000 |
| 8 | 10.0000 | 0.13572 | 16 | 10.0000 | 1.00000 |

**Table 7. Kinetic parameters to be estimated in the HIV model**

| Exp. | $I_0$ (μM) | Parameter | Search Space | Exp. | $I_0$ (μM) | Parameter | Search Space |
|---|---|---|---|---|---|---|---|
| - | - | *ks* ($s^{-1}$) | $10^{-6} – 10^{6}$ | A | 0.0000 | $E_0$ (μM) | 0.002 – 0.006 |
| - | - | *kcat* ($s^{-1}$) | $10^{-6} – 10^{6}$ | B | 0.0015 | $E_0$ (μM) | 0.002 – 0.006 |
| - | - | *kp* ($s^{-1}$) | $10^{-6} – 10^{6}$ | C | 0.0030 | $E_0$ (μM) | 0.002 – 0.006 |
| - | - | *ki* ($s^{-1}$) | $10^{-6} – 10^{6}$ | D | 0.0040 | $E_0$ (μM) | 0.002 – 0.006 |
| - | - | *kde* ($s^{-1}$) | $10^{-6} – 10^{6}$ | E | 0.0040 | $E_0$ (μM) | 0.002 – 0.006 |
| A | 0.0000 | $S_0$ (μM) | 10 – 40 | A | 0.0000 | *offset* | -0.100 – 0.100 |
| B | 0.0015 | $S_0$ (μM) | 10 – 40 | B | 0.0015 | *offset* | -0.100 – 0.100 |
| C | 0.0030 | $S_0$ (μM) | 10 – 40 | C | 0.0030 | *offset* | -0.100 – 0.100 |
| D | 0.0040 | $S_0$ (μM) | 10 – 40 | D | 0.0040 | *offset* | -0.100 – 0.100 |
| E | 0.0040 | $S_0$ (μM) | 10 – 40 | E | 0.0040 | *offset* | -0.100 – 0.100 |

### Figures

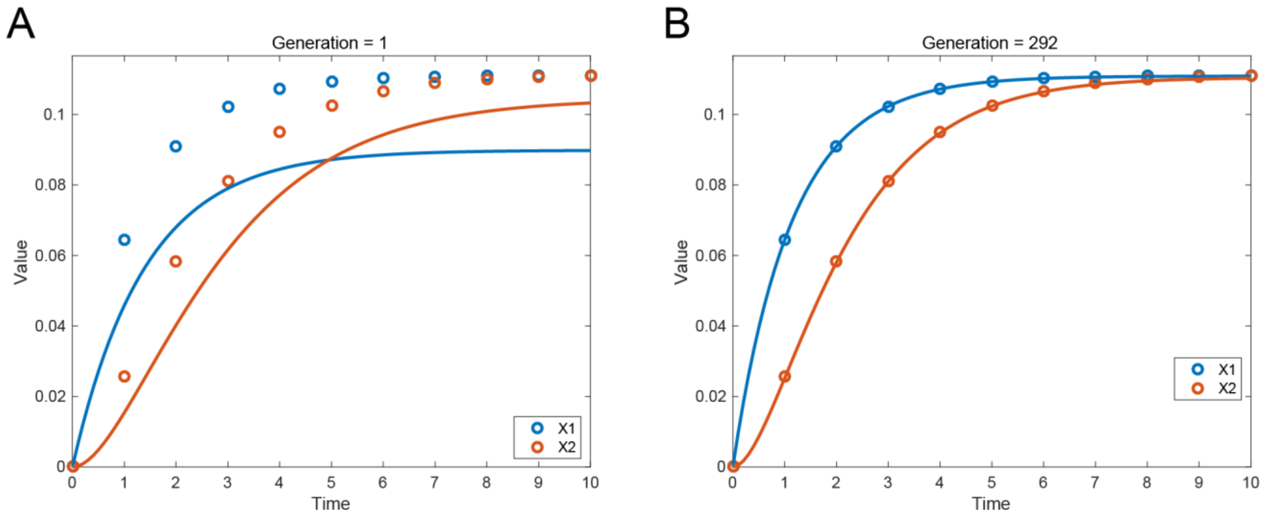

**Figure 1. Examples of figures that appear during parameter estimation**

A figure pops up just after starting **RCGA_UNDXMGG_PE** or **RCGA_REXstarJGG_PE**, and it will be updated during parameter estimation. The circles and the lines indicate the target experimental data and the current best model behavior, respectively. (A) The behavior of the best parameter set at the 1$^{st}$ generation. (B) The behavior of the best parameter set at the 292$^{nd}$ generation.

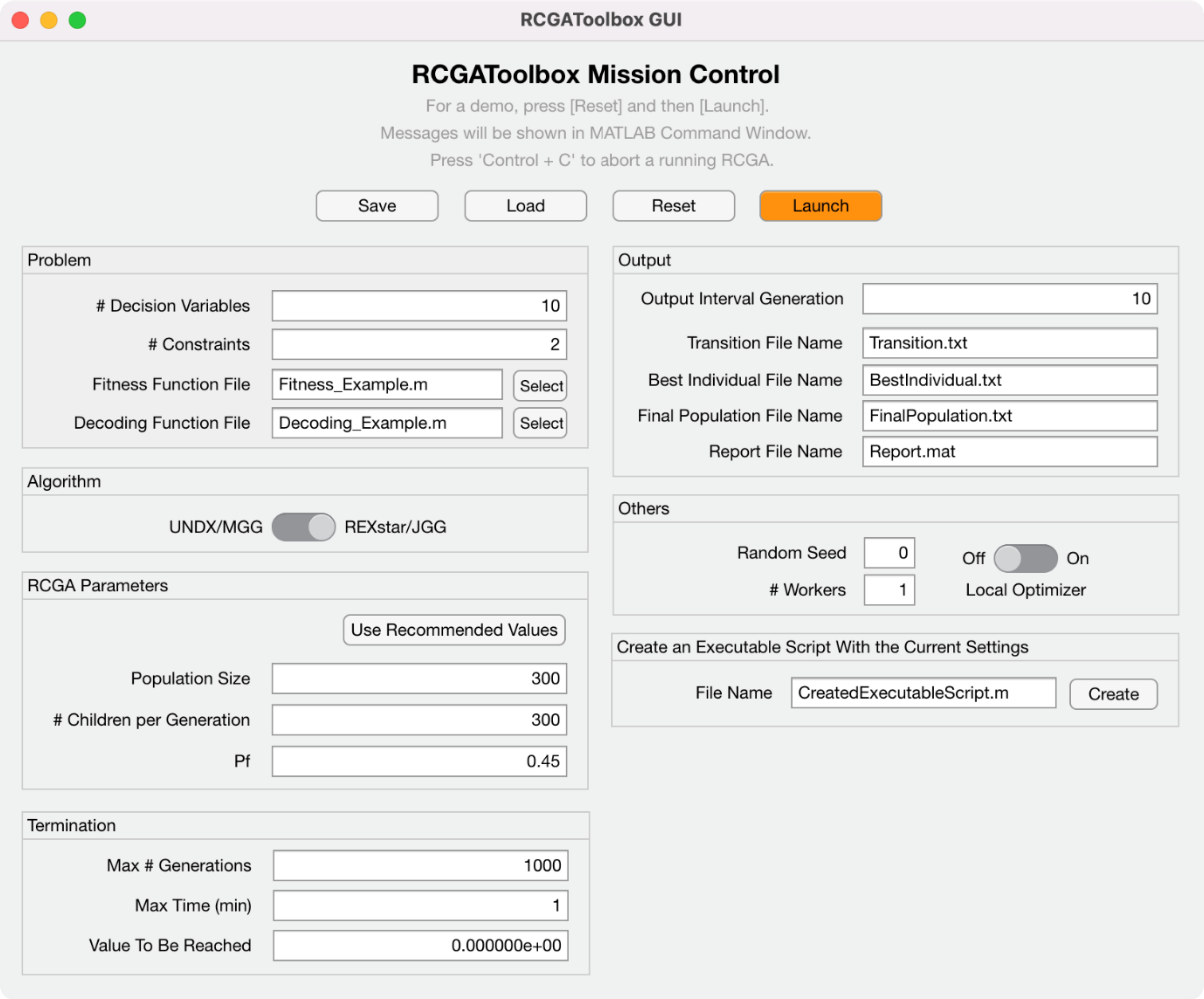

**Figure 2. GUI for general optimization problems**

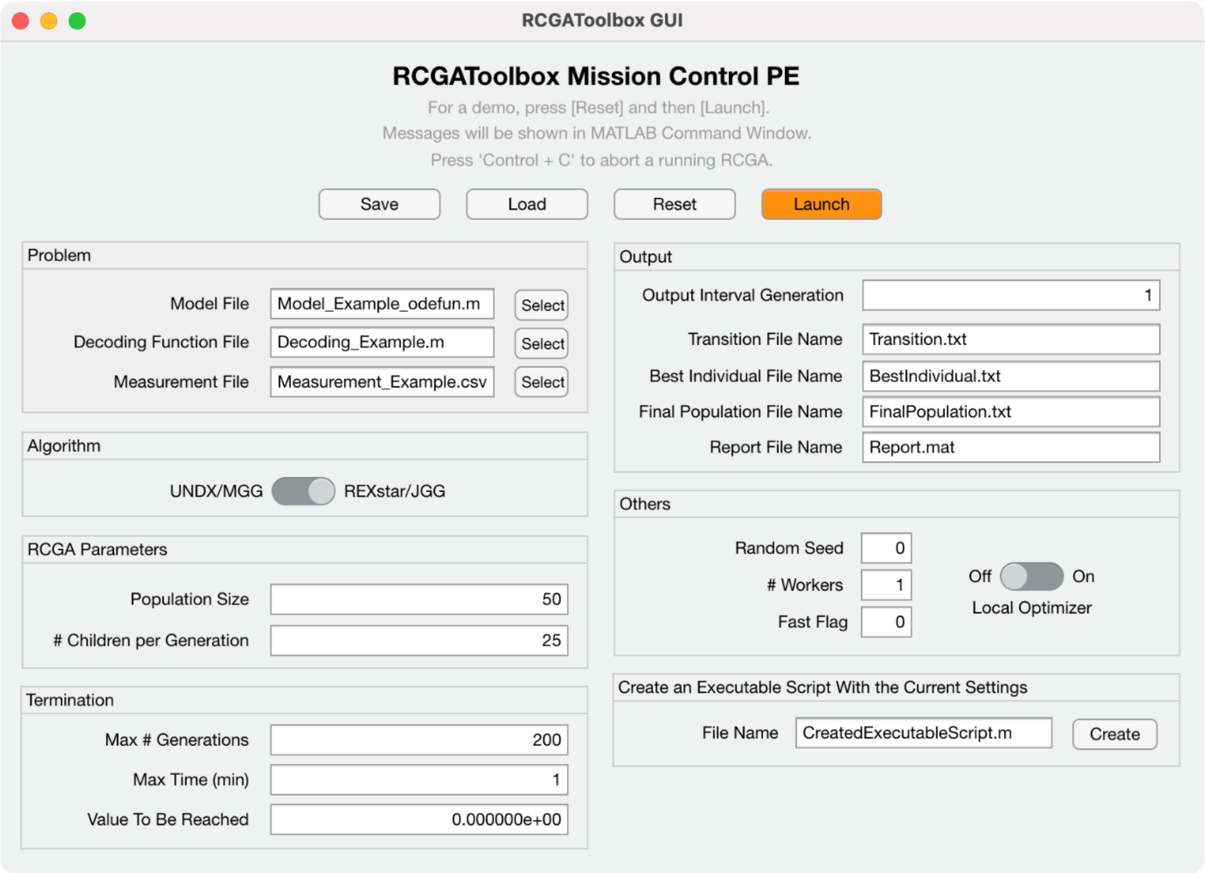

**Figure 3. GUI for parameter estimation in systems biology**

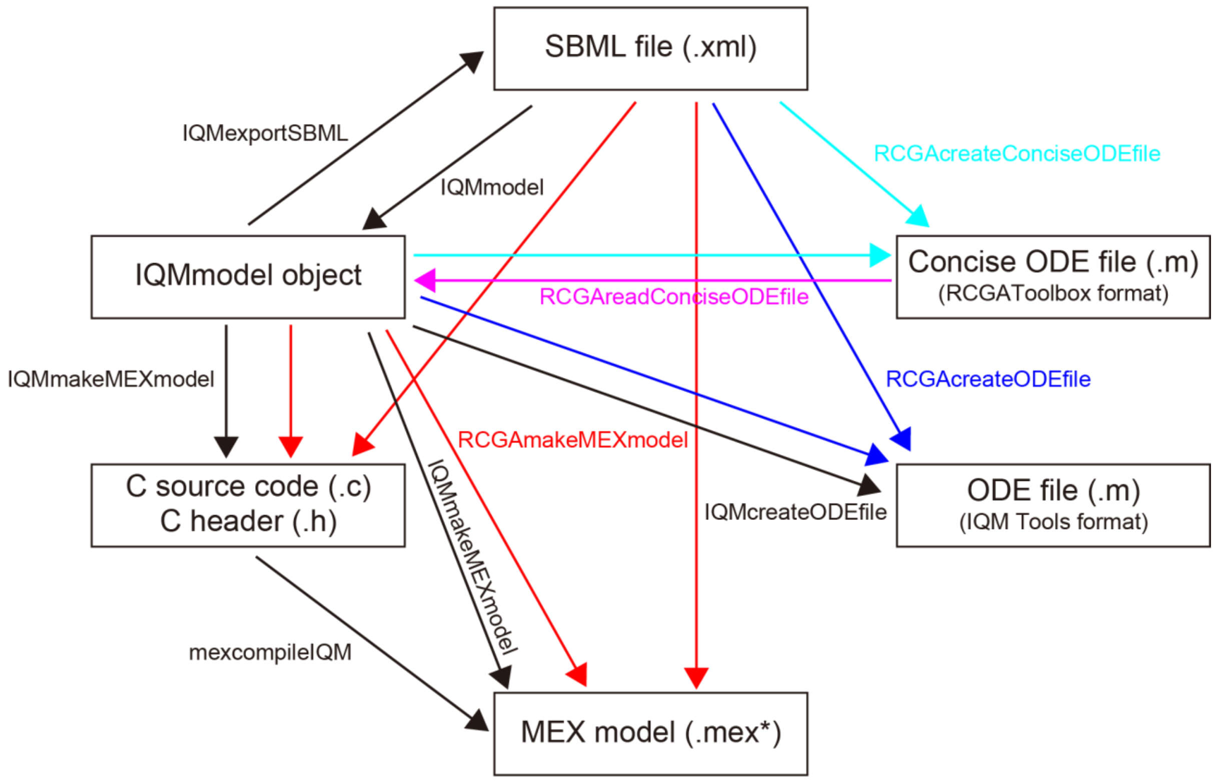

**Figure 4. Model formats and conversion functions**

The rectangles and arrows indicate the model formats and conversion functions, respectively. For example, the function RCGAreadConciseODEfile converts a concise ODE file into an IQM model object. The conversion functions highlighted in red, blue, cyan, and magenta are provided by RCGAToolbox. Those indicated by black arrows are provided by IQM Tools.

```
for i = 1 to n_ps do
    for j = 1 to n_ps − 1 do
        Sample u ∈ U(0,1)
        if [φ(x_j) = φ(x_{j+1})] or [u < P_f] then
            if [f(x_j) > f(x_{j+1})] then
                swap(x_j, x_{j+1})
            fi
        else
            if [φ(x_j) > φ(x_{j+1})] then
                swap(x_j, x_{j+1})
            fi
        fi
    od
    if no swap done
        break
    fi
od
```

**Figure 5. Stochastic ranking using a bubble-sort-like procedure**

$x_j$ indicates the $j^{th}$-ranked individual (i.e., decision variable vector). The initial ranking is generated randomly. $U(0, 1)$ is a uniform random number generator, and $n_{ps}$ is the population size. $P_f$ specifies the probability that only the objective function $f$ is used to compare individuals when they have different values of the penalty function $\phi$. It has been demonstrated that the stochastic ranking works well when $0.4 < P_f < 0.5$ (Runarsson and Yao, 2000).

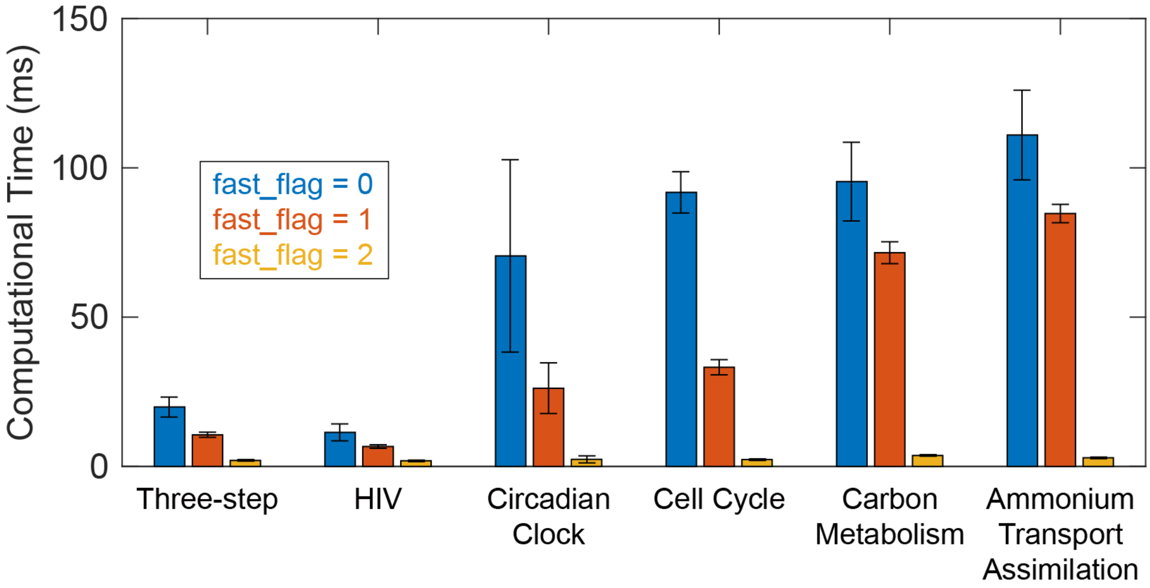

**Figure 6. Computational times of different simulation methods**

Results are expressed as mean ± SD ($n$ = 100). In the case of **fast_flag** = 0, the MATLAB built-in ODE solver ode15s was used with the option BDF = 'on'. In the case of **fast_flag** = 1, a pre-compiled ODE solver (CVODE) provided by SundialsTB was used. In the case of **fast_flag** = 2, the models were compiled using IQM Tools and combined with a pre-compiled ODE solver (CVODE). In all three cases, the same numerical integration algorithm (BDF: backward differentiation formula) was internally used. The kinetic models for three-step and HIV are described in **Appendix E**. The kinetic models for the circadian clock (BIOMD0000000016), cell cycle (BIOMD0000000005), carbon metabolism (BIOMD0000000051), and ammonium transport assimilation (MODEL1901090001) were obtained from BioModels (Malik-Sheriff, et al., 2020). For all experiments, both the absolute and relative tolerances were set to $10^{-6}$. The computational experiment was performed on a single core of Intel Xeon E5-1650 v4 with Windows 10 (2004) using MATLAB R2016a.

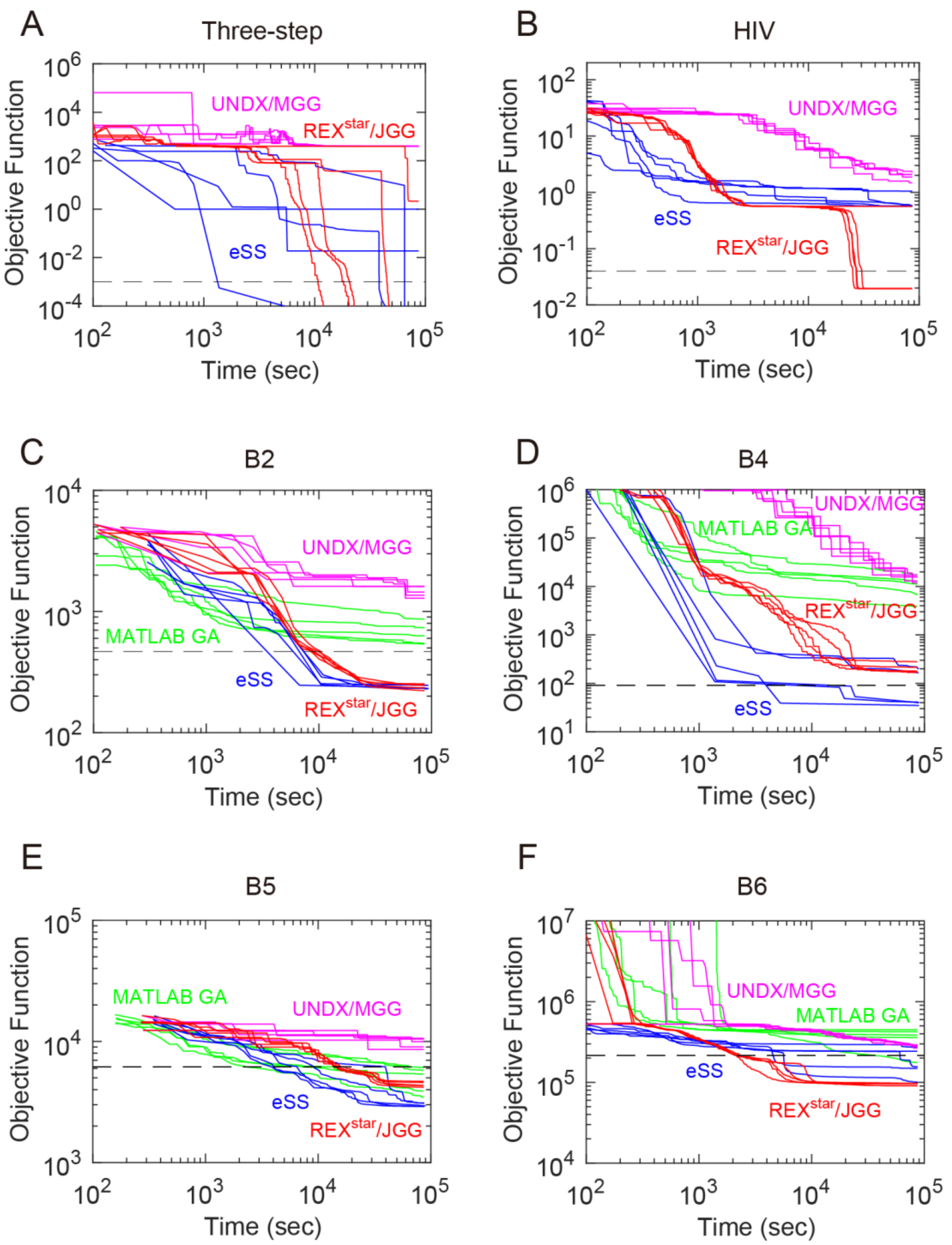

**Figure 7. Convergence curves for biological test problems**

Five independent runs were carried out for each algorithm. We employed MATLAB GA (Global Optimization Toolbox, MathWorks, Inc) and eSS [R2019 from MEIGO (Egea, et al., 2014)] as controls. We used RCGAToolbox for UNDX/MGG and REX$^{star}$/JGG. Black dashed lines show the values to be reached (VTRs). In (A) and (B), the convergence curves for MATLAB GA are not shown in the figures as MATLAB GA froze at the first generation without generating any output. The computational experiment was performed on a single core of Intel Xeon E5-4650 v3 with SUSE Linux Enterprise Server 11 (x86_64) using MATLAB R2016a.

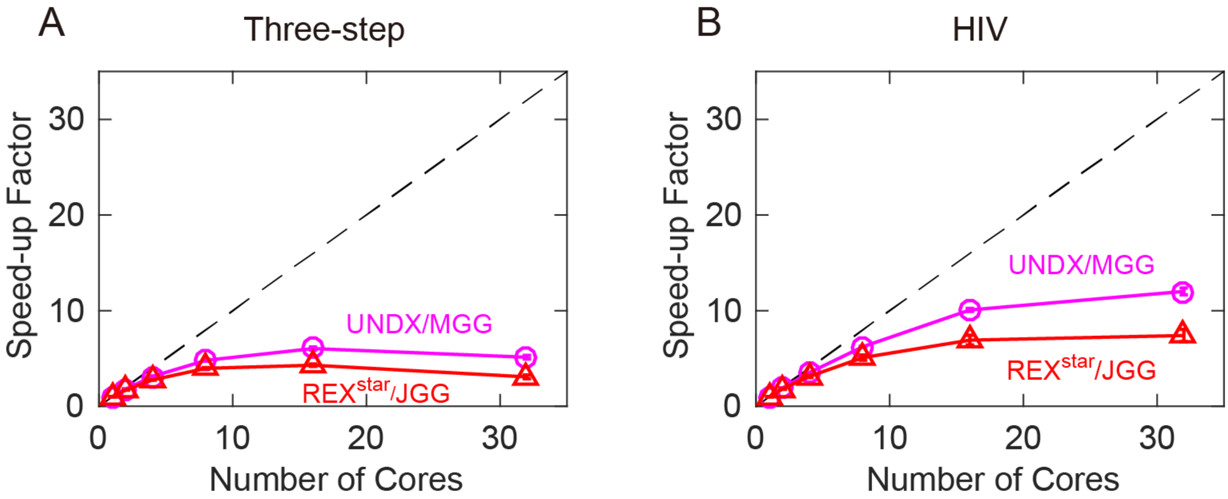

**Figure 8. Speed-up by parallel computation**

(A) The three-step problem. (B) The HIV problem. Five independent runs were carried out. Circles and triangles represent mean values ($n$ = 5). Error bars represent ± SD (the error bars are very small in the figures). The black dashed diagonal lines show ideal linear speed-up (i.e. the calculation speed is doubled if the number of cores is doubled). Speed-up factor is the ratio between the computational speed for parallel computation and for sequential computation. The number of cores indicates the computation cores used for parallel computation. We used RCGAToolbox for UNDX/MGG and REX$^{star}$/JGG with the local optimizer and the stochastic ranking ($P_f$ = 0.45). The computational experiment was performed on Intel Xeon E5-4650 v3 with SUSE Linux Enterprise Server 11 (x86_64) using MATLAB R2016a.

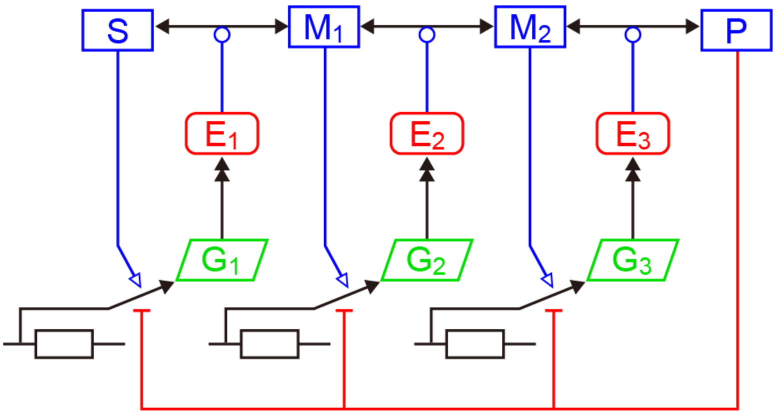

**Figure 9. Network map of the three-step model**

*S* and *P* are the substrate and the product, respectively. $G_i$, $E_i$, and $M_i$ are the $i^{th}$ mRNA, enzyme, and intermediate, respectively. For simplicity, CADLIVE notation was used (Kurata, et al., 2007; Kurata, et al., 2005; Kurata, et al., 2003): transcription ( ), translation ( ), enzymatic catalysis ( ), reversible reaction ( ), activation ( ), and repression ( ).

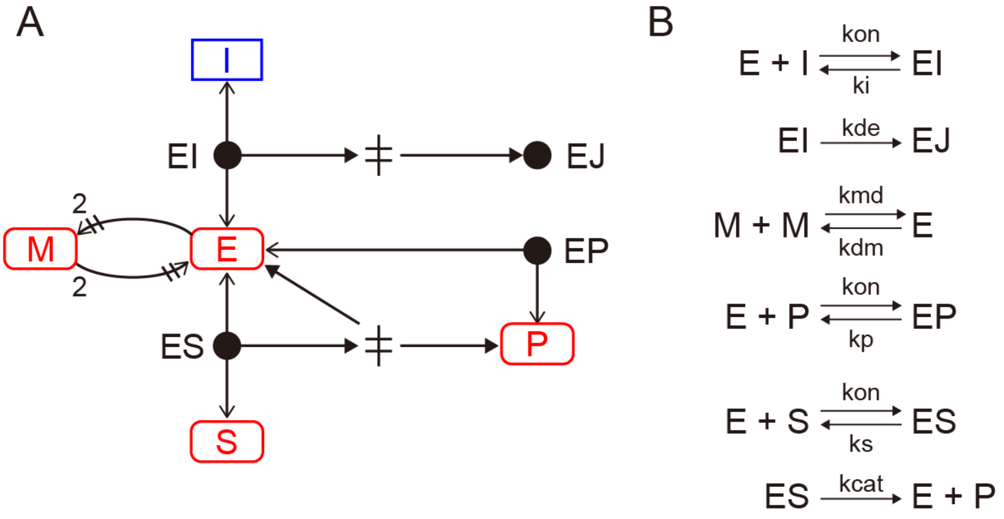

**Figure 10. HIV model**

(A) Network map. CADLIVE notation was used for simplicity (Kurata, et al., 2007; Kurata, et al., 2005; Kurata, et al., 2003): Binding (←●→), irreversible reaction (⟶), homo association and dissociation (⟶⫲), and transition (ǂ). (B) Reaction equations. The enzyme is inactive in a monomer form *M* and active in a dimer form *E*. The product *P* is a competitive inhibitor for the substrate *S*. *I* is the irreversible inhibitor. The different complexes are the enzyme-inhibitor (*EI*), irreversibly inactivated enzyme-inhibitor (*EJ*), enzyme-product (*EP*), and enzyme-substrate (*ES*) complex, respectively.
